# The mechanism of mammalian proton-coupled peptide transporters

**DOI:** 10.1101/2024.02.04.578827

**Authors:** Simon M Lichtinger, Joanne L Parker, Simon Newstead, Philip C Biggin

## Abstract

Proton-coupled oligopeptide transporters (POTs) are of great pharmaceutical interest owing to their promiscuous substrate binding site that has been linked to improved oral bioavailability of several classes of drugs. Members of the POT family are conserved across all phylogenetic kingdoms and function by coupling peptide uptake to the proton electrochemical gradient. Cryo-EM structures and alphafold models have recently provided new insights into different conformational states of two mammalian POTs, SLC15A1 and SLC15A2. Nevertheless, these studies leave open important questions regarding the mechanism of proton and substrate coupling, while simultaneously providing a unique opportunity to investigate these processes using molecular dynamics (MD) simulations. Here, we employ extensive unbiased and enhanced-sampling MD to map out the full SLC15A2 conformational cycle and its thermodynamic driving forces. By computing conformational free energy landscapes in different protonation states and in the absence or presence of peptide substrate, we identify a likely sequence of intermediate protonation steps that drive inward-directed alternating access. These simulations identify key differences in the extracellular gate between mammalian and bacterial POTs, which we validate experimentally in cell-based transport assays. Our results from constant-PH MD and absolute binding free energy (ABFE) calculations also establish a mechanistic link between proton binding and peptide recognition, revealing key details underpining secondary active transport in POTs. This study provides a vital step forward in understanding proton-coupled peptide and drug transport in mammals and pave the way to integrate knowledge of solute carrier structural biology with enhanced drug design to target tissue and organ bioavailability.

## Introduction

Cells require an external lipid membrane to separate their internal cytoplasm from the environment. Since the membrane permeability of common solutes spans ten orders of magnitude, some molecules diffuse readily across the membrane, while the translocation of others requires facilitation by carriers. (***Stillwell, 2016***; ***Pizzagalli et al., 2021***) Understanding the processes by which small molecules cross membranes is of key pharmacological interest owing to their role in drug delivery, which may be mediated by passive diffusion, protein carriers, or a combination of both. (***Sugano et al., 2010***) The solute-carrier (SLC) superfamily encompasses 65 families of more than 450 genes, with substrates ranging in size simple ions to complex macromolecules used in metabolism and signalling. (***Pizzagalli et al., 2021***) Within this superfamily, the SLC15 family includes proton-coupled oligopeptide transporters (POTs) that have significant homology through all domains of life and are evolutionarily ancient. (***Daniel et al., 2006***) Of the four mammalian family members, PepT1 (SLC15A1) and PepT2 (SLC15A2) are the most well studied. The former is predominantly expressed in the small intestine and characterised as a low-affinity, high-capacity transporter. (***Fei et al., 1994***) The latter has a broader expression pattern including the kidneys, lungs and brain and is described as high-affinity, low-capacity. (***Daniel and Kottra, 2004***) As secondary-active transporters, they couple uphill substrate translocation to the symport of protons down their electrochemical gradient. (***Fei et al., 1994***; ***Rubio-Aliaga et al., 2000***) The peptide–proton stoichiometry is not conserved between different substrates and POT family members. (***Parker et al., 2014***) For PepT1, stoichiometries of 1:1 and 2:1 have been reported for neutral / basic and acidic di-peptides, respectively. (***Fei et al., 1994***; ***Steel et al., 1997***) For PepT2, a 2:1 stoichiometry was reported for the neutral di-peptide D-Phe-L-Ala and 3:1 for anionic D-Phe-L-Glu. (***Chen et al., 1999***) Alternatively, ***Fei et al. (1999)*** have found 1:1 stoichiometries for either of D-Phe-L-Gln (neutral), D-Phe-L-Glu (anionic), and D-Phe-L-Lys (cationic). Here, we work under the assumption of a 2:1 stoichiometry for neutral di-peptides, motivated also by our computational results that indicate distinct and additive roles played by two protons in the conformational cycle mechanism.

POT family transporters belong to the major facilitator superfamily (MFS) and share a conserved topology of two six helix bundles that form the functional transport domain, their N- and C-termini facing the cytoplasm. (***Newstead et al., 2011***) They operate via an alternating access mechanism encoded in four inverted topology repeats, progressively reorienting the N- and C-terminal bundles to cycle through outwards-facing (OF), occluded (OCC) and inwards-facing (IF) states. (***Radestock and Forrest, 2011***) Since the first structure of a POT family member was published (***Newstead et al., 2011***), many procaryotic (***Solcan et al., 2012***; ***Guettou et al., 2013***; ***Doki et al., 2013***; ***Lyons et al., 2014***; ***Guettou et al., 2014***; ***Zhao et al., 2014***; ***Fowler et al., 2015***; ***Boggavarapu et al., 2015***; ***Beale et al., 2015***; ***Parker et al., 2017***; Martinez Molledo et al., 2018; ***Ural-Blimke et al., 2019***; ***Minhas and Newstead, 2019***; ***Stauffer et al., 2022***; ***Kotov et al., 2023***) and plant (***Parker and Newstead, 2014***; ***Sun et al., 2014***) homologues have been structurally and biochemically characterised, all in IF states with varying degrees of occlusion (see figure 1a for an overview of available POT structures and their conformational states). Several residues have been suggetsed to be involved in proton transfer, including a partially conserved histidine on TM2 (H87; residue numbers refer to PepT2, if not specified otherwise) (***Terada et al., 1996***; ***Fei et al., 1997***; ***Chen et al., 2000***; ***Omori et al., 2021***; ***Parker et al., 2021***) and two conserved glutamates on TM1 (E53 and E56) (***Jensen et al., 2012***; ***Doki et al., 2013***; ***Aduri et al., 2015***), while simulations have helped our understanding of proton-transfer processes and conformational changes. (***Parker et al., 2017***; ***Selvam et al., 2018***; ***Batista et al., 2019***; ***Li et al., 2022***) However, the details of the molecular mechanism of alternating access in POTs — particularly regarding the coupling of conformational changes, substrate binding and proton movement to each other — remain unclear.

**Figure 1.**
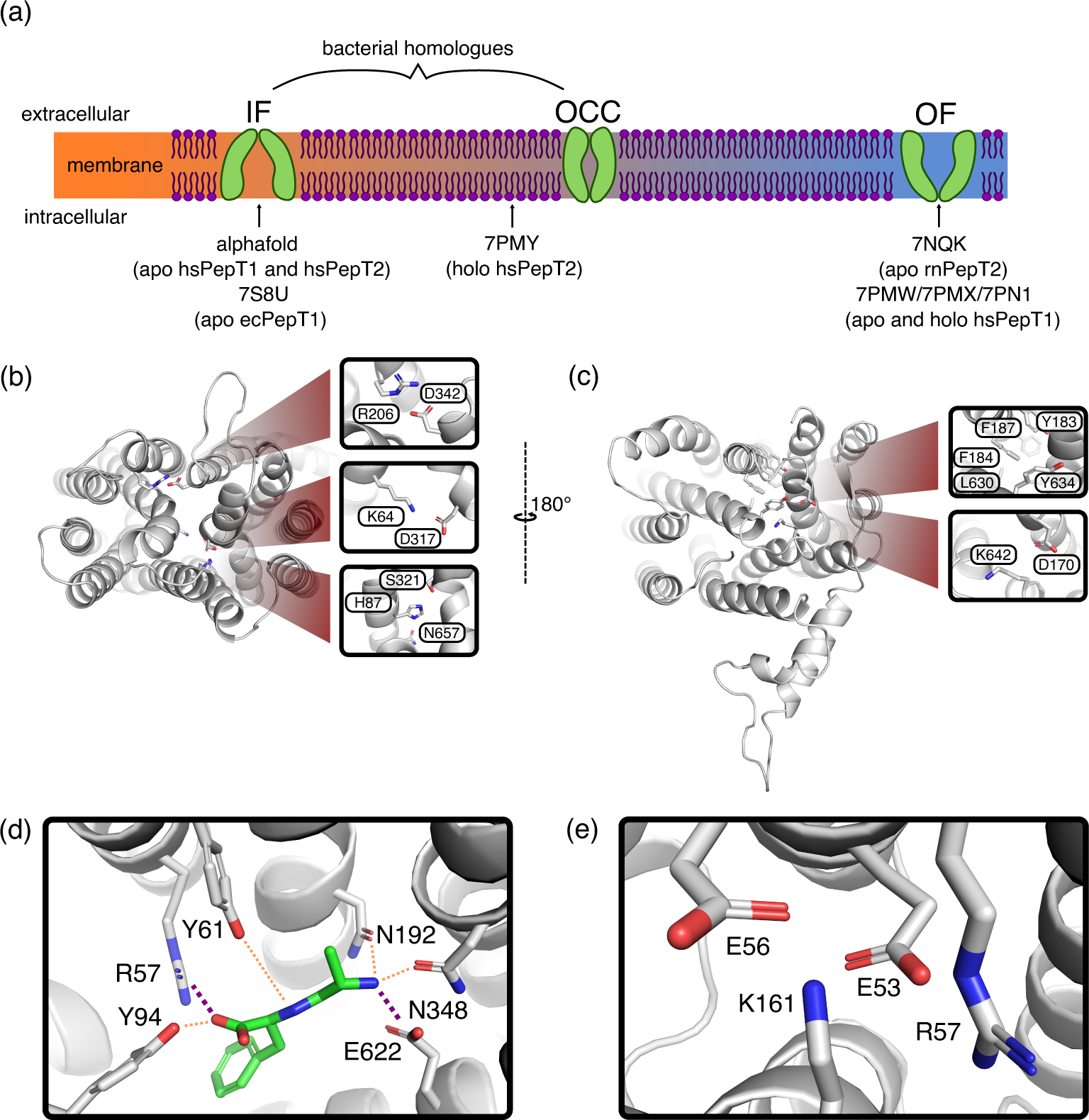
(a) Schematic overview of the conformational diversity of available mammalian POT structures. Intermediate positions between states indicate partial gate opening. (b) Alphafold-predicted inwards-facing (IF) HsPepT2 structure (top view), highlighting potential inter-bundle extracellular gate interactions. (c) Outwards-facing (OF) Cryo-EM structure of apo RnPepT2 (7NQK, bottom view) (***Parker et al., 2021***), highlighting potential inter-bundle intracellular gate interations. (d) Ala-Phe substrate binding pose, representative cluster frame of 1 µs MD simulation from 7NQK structure with added ligand, for setup details see Methods and Materials. Purple dotted lines represent salt-bridge contacts, orange dotted lines other polar contacts. (e) ExxER motif salt-bridge cluster, representative cluster frame of 1 µs MD simulation from 7NQK structure.

Cryo-EM and Alphafold 2 have recently provided views of mammalian POTs in conformations spanning from OF via inward-facing, partially occluded to fully-open IF. (***Parker et al., 2021***; ***Killer et al., 2021***; ***Shen et al., 2022***; ***Jumper et al., 2021***) From these structures emerges a picture where the intracellular gate is constituted by broad close-packing of hydrophobic residues on TM 4, 5, 10 and 11, with possible stabilisation from the conserved D170–K642 salt-bridge. The extracellular gate appears to be spread along the cleft between the N- and C-terminal bundles, with contributions from the H87 (TM 2) – S321 (TM 7) polar interaction network as well as the R206 (TM 5) – D342 (TM 8) and K64 (TM 1) – D317 (TM 7) salt bridges (figure 1b). This is intriguing, because the mammalian H87 residue is only conserved in some prokaryotic homologues, and R206–D342 just among mammalian POTs. We speculate based on this feature that the extracellular gating mechanism could be less conserved than POT alternating access in general. As for the intracellular gating mechanism, an involvement of the D170–K642 saltbridge has been suggested, and the OF structure shows close-packing of several hydrophobic residues (F184, Y183, F187, L630 and Y634) that constrict access to the binding site from the intracellular side (figure 1c). (***Parker et al., 2021***) It is not known thus far how the opening of the intracellular gate (i.e., the OCC→IF transition) is triggered, and how it is coupled to proton movement and the presence of substrate.

POTs accommodate their substrates in a highly conserved binding pocket, interfacing between an acidic patch on the C-terminal bundle and a basic patch on the N-terminal bundle (figure 1c). For di-peptides, the N-terminus is coordinated by E622 (TM 10) together with N192 (TM 5) and N348 (TM 8), while the C-terminus engages R57 (TM 1, or the equivalent lysine residue in PepT1) as well as Y94 (TM 2). Another conserved tyrosine, Y61 (TM 1), hydrogen-bonds to features of the peptide backbone. (***Lyons et al., 2014***; Martinez Molledo et al., 2018; ***Killer et al., 2021***) Tri-peptides may adopt a similar orientation as di-peptides (***Guettou et al., 2014***), or sit vertically in the transporter binding pocket (***Lyons et al., 2014***), although it has been suggested that this vertical electron density could alternatively be explained by a bound HEPES molecule. (Martinez Molledo et al., 2018) Considering the consensus structural interaction pattern, we decided to investigate primarily the role of E622 and R57 in holding the substrate, and also note that R57 is part of the highly conserved E^53^xxER motif (figure 1d). (***Newstead, 2017***) The second glutamate in this motif (E56) in particular has been linked to proton coupling experimentally. (***Jensen et al., 2012***) Since R57 interacts with both the ExxER glutamates and the substrate C-terminus, we hypothesise that it may play an important role in substrate–proton coupling.

In this study we use extensive unbiased and enhanced-sampling molecular dynamics (MD) simulations (totalling close to 1 ms of sampling) to show how changes in protonation states of H87 and D342 control the OCC↔OF transition as an extracellular gate. We validate the importance of these residues for transporter function in cell-based transport assays. We also elucidate the role of E^53^xxER glutamates and the substrate-engaging E622 in controlling the OCC↔IF transition, thereby identifying a clear molecular basis for the directionality of proton movement coupling to conformational changes. Furthermore, we establish several distinct effects of the presence of substrate, coupling ligand binding with protein conformational changes and also linking it to protonation of the E56 and E622 titratable residues. Taken together, our work provides for the first time a detailed model of a plausible sequence of steps for substrate and proton-coupled alternating access in mammalian POTs.

## Results

### Unbiased MD identifies extra- and intra-cellular gate opening triggers

We began our computational investigation by embedding PepT2 structures (using the sequence of the rat homologue) in the OF (***Parker et al., 2021***), IF (alphafold prediction, ***Jumper et al. (2021)***) and inwards-facing partially occluded (***Killer et al., 2021***) conformations in 3:1 POPE:POPG membranes (figure S1, details in Materials and Methods). While we were able to obtain stable wide-open OF and IF simulation boxes, the OCC state required further attention as the MD simulations from the inwards-facing, partially-occluded structure showed embedding artifacts, including extracellular-gate instability and intracellular gate hydrophobic collapse (figure S1a). We therefore opted to derive an OCC state using metadynamics simulations in 5 replicates (figure S2, further details in Materials and Methods), using stability in unbiased MD to select the best among the obtained candidates. The OCC state thus developed is validated by the further work in this paper, showing it to be a stable conformational basin that is functionally occluded in that it can open both towards OF and IF in different protonation state conditions (figures 2, 3 and 4).

**Figure 2.**
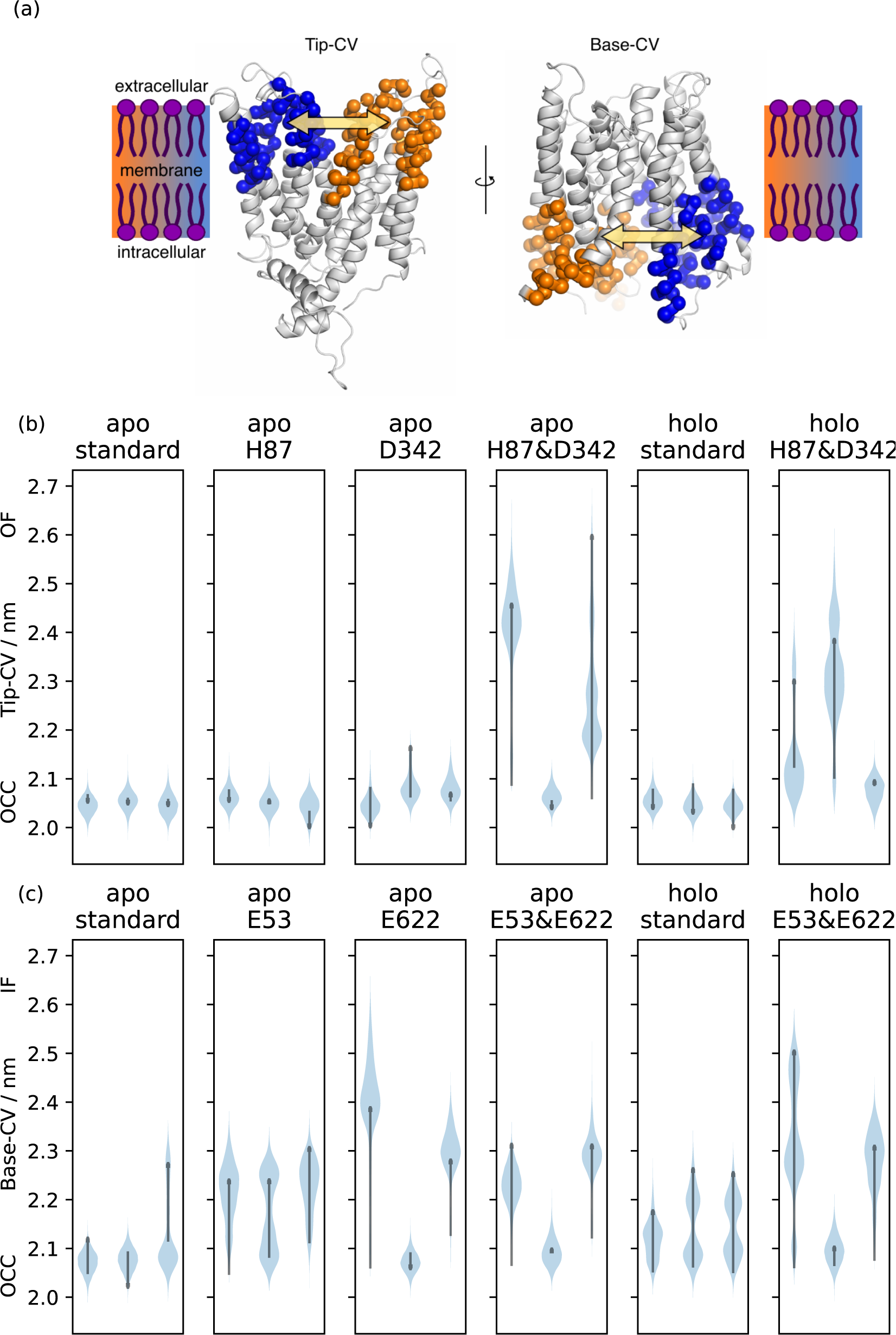
(a) Illustration of the CVs used to quantify extra- and intracellular gate opening, consisting of inter-bundle centre-of-mass distances between the helical tips (top 11 residues) and bases (bottom 11 residues). (b)+(c) Triplicate 1 µs-MD simulations starting from OCC, showing the effects of different protonation and substrate binding states, projected onto the (b) Tip-CV and (c) Base-CV respectively. Violin plots are trajectory histograms, arrows link the CV values of the first and last frames.

**Figure 3.**
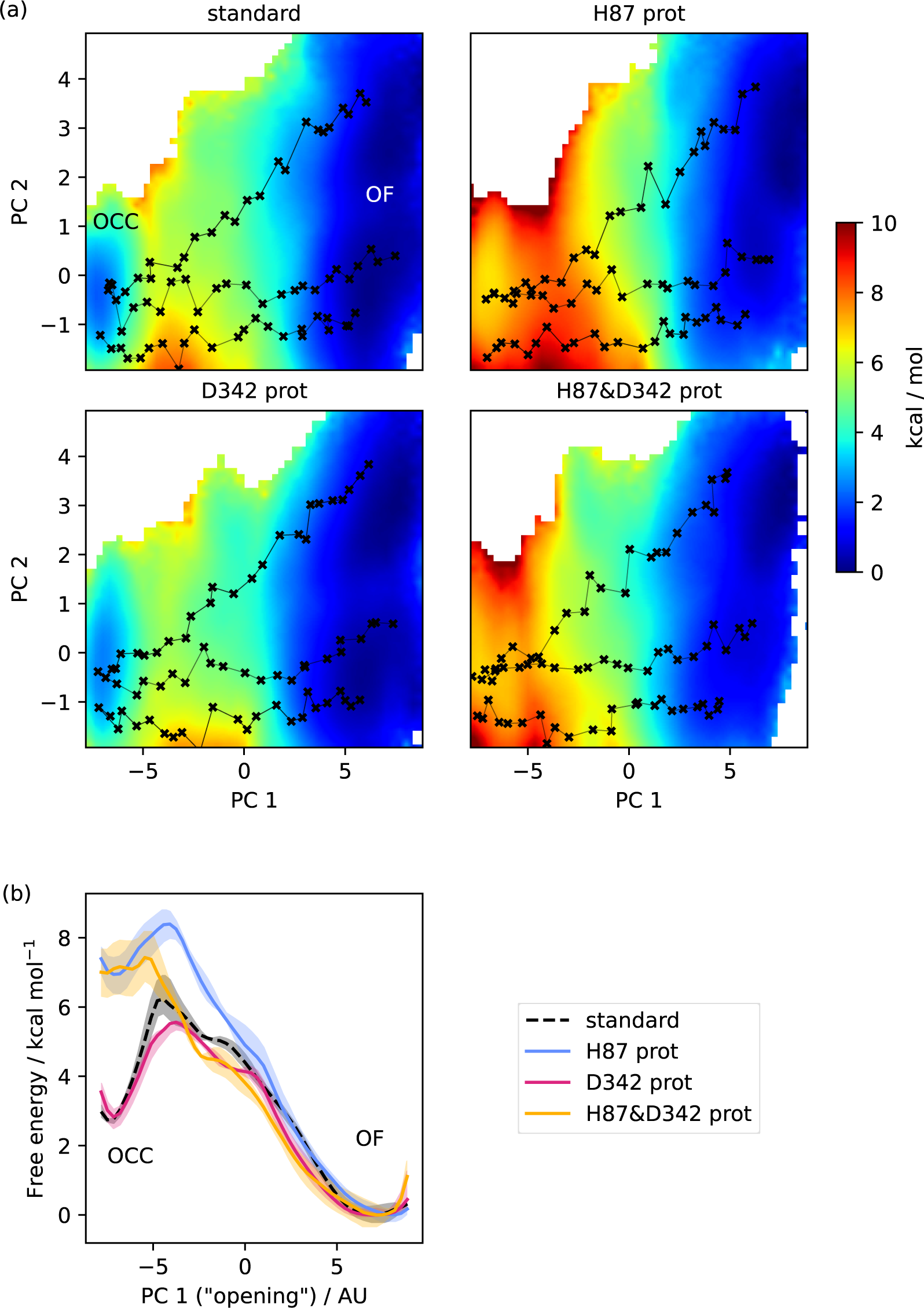
(a) 2D-PMFs from REUS starting with MEMENTO paths, in different protonation states of candidate extracellular gating residues. (b) Projection of the 2D-PMFs in part a onto PC 1 using Boltzman reweighting. Shaded areas indicate convergence errors as the range of PMF values for a given CV value obtained with the first 40 %, the last 40 % and 100 % of sampling included (after alignment to the 100 % curve). H87 and D342 form an additive extracellular gate, where H87 protonation changes the relative OCC–OF state energies as well as the transition barrier, while D342 protonation only contributes in the transition region. Note that the individual PMFs are only determined by our REUS approach up to additive constants, and are shown aligned here at the OF state for convenience of comparison.

**Figure 4.**
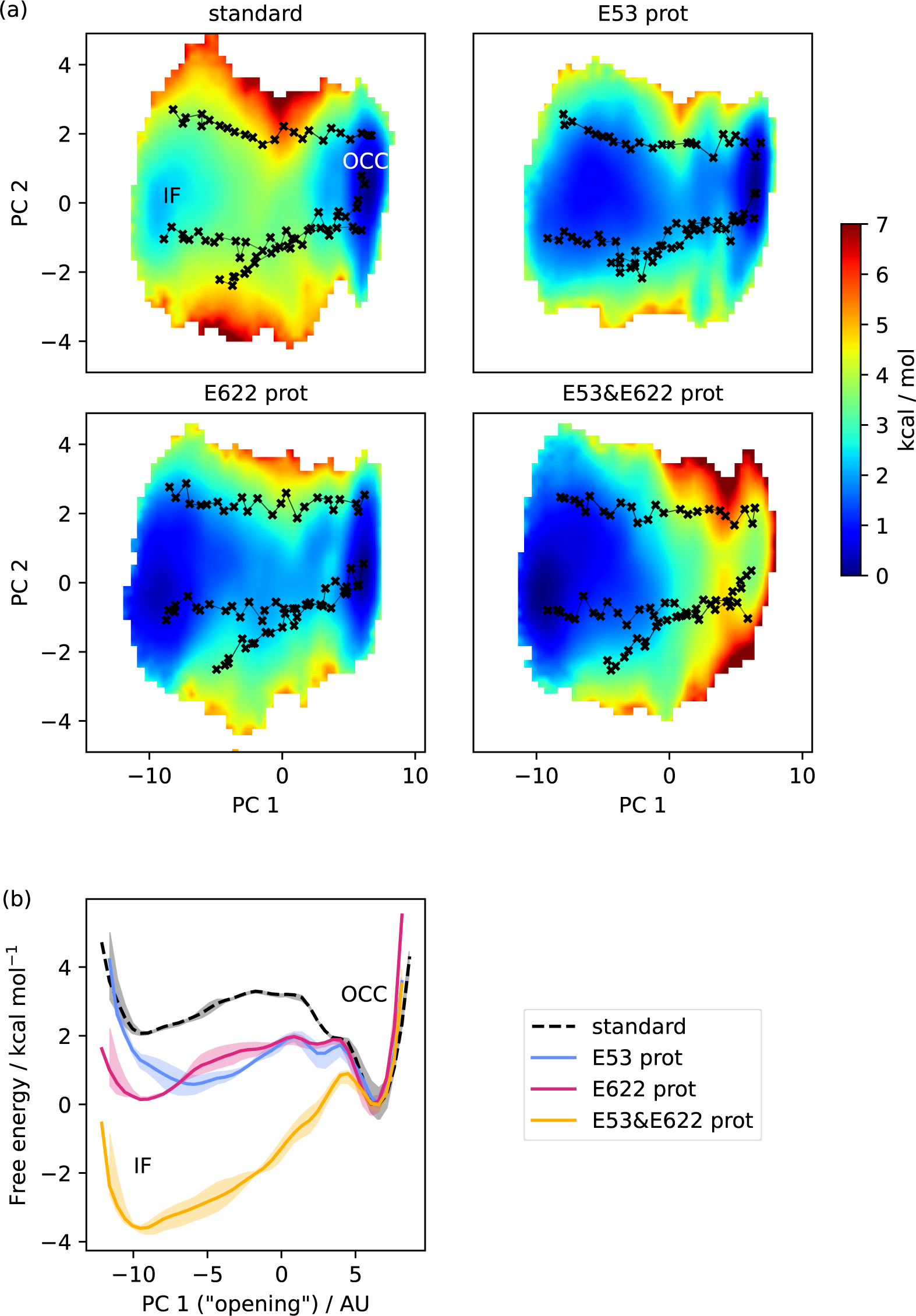
(a) 2D-PMFs from REUS starting with MEMENTO paths, in different protonation states of candidate intracellular gate-controlling residues. (b) Projection of the 2D-PMFs in part a onto PC 1 using Boltzman reweighting. Shaded areas indicate convergence errors as the range of PMF values for a given CV value obtained with the first 40 %, the last 40 % and 100 % of sampling included (after alignment to the 100 % curve). E53 and E622 protonation have additive and approximately equal effects on driving the OCC→IF transition. Note that the individual PMFs are only determined by our REUS approach up to additive constants, and are shown aligned here at the IF state for convenience of comparison.

Equipped with models of the protein conformations required for PepT2 alternating access (OF, OCC and IF), we ran triplicate sets of 1 µs-long MD simulations in a range of conditions. To decide which conditions to probe apart from the apo, standard protonation states as obtained above, we investigated the extent to which the H87–S321, R206–D342, K64–D317 and D170–K642 interactions (see figure 1b–c) are formed in the OF (closed intracellular gate) and IF (closed extracellular gate) conformational states (figure S3). In the IF state, we found that H87–S321 and D342–R206 are consistently interacting, whereas the K64–D317 interaction, while formed in ≈ 72% of MD frames, is unstable and of a transient nature and therefore unlikely to contribute much to extracellular gate stability. The D170–K642 saltbridge, in turn, is only formed in ≈ 22% of OF-state frames, thus likely not substantially adding to the stability of the intracellular gate. We therefore decided to mainly focus on probing the H87–S321 and D342–R206 interactions with respect to control of the extracellular gate. Since no saltbridges or other specific interactions involving protonatable residues seem to demarcate the intracellular gate, we decided to focus on the ExxER glutamates and E622 for their effects on intracellular gate opening.

Guided thus in our choice of which residues to investigate, we probed whether the OCC state opens spontaneously to OF or IF states in a range of different protonation-state and mutation conditions (as assessed by projection of unbiased MD runs onto intuitive collective variables, or CVs, defined as the centre-of-mass distance between the tips and bases of the N- and C-terminal bundles, see figure 2a). We found that the extracellular gate remains stably closed in triplicates of 1 µs-long MD when H87 or D342 are protonated individually, but the OCC state can open spontaneously on the simulated time scale to an OF conformation when both are protonated simultaneously (figure 2b; figure S4a for plots of the opposite gate in the same trajectories, showing how flexibility of the intra- and extracellular gates is anti-correlated). A comparable effect is found in the presence of the physiogical peptide substrate L-Ala–L-Phe (figure 2b, panels 5–6). We have also tested further combinations of mutations and protonation state changes relating to the putative extracellular gating interactions (D317 protonation and mutations of R206, S321, D342 to alanine, with and without H87 protonation) as well as some control mutants which we did not expected to have an effect (I135L, T202A, Q340A and the saltbridge-swapped mutant R206D & D342R, combined with H87 protonation). Across our 48 * 1 µs unbiased MD runs collated in figure S5, we observed 3 full extracellular-gate opening events, in conditions where H87 was protonated and the D342–R206 saltbridge was also disrupted either by D342 protonation or mutation to alanine (D342A). We also saw one partial opening event when in addition to H87 protonation we also mutated S321 to alanine (S321A). The data thus suggests that for spontaneous extracellular-gate opening to occur on this time scale in unbiased MD, disruption of the OCC-state H87 interaction network is essential, and the D342 salt bridge appears to make an additive contribution towards extracellular-gate stability (though this is not a strict correlation, as illustrated by the S321A partial opening event). The intracellular gate, by contrast, is more flexible than the extracellular gate even in the apo, standard protonation state; however, following either protonation of the conserved E53 and E622 residues or the insertion of Ala-Phe substrate, the intracellular gate becomes more flexible and can spontaneously open (figure 2c; see figure S4b for the corresponding plots of the extracellular gate opening).

Although these unbiased simulations show a large amount of stochasticity and drawing clean conclusions from the data is difficult, we can already appreciate a possible mechanism where protons move down the transporter pore, first engaging H87 and D342 to favour the OF state and then moving to ExxER and E622 to favour the IF orientation, driving successive conformational changes along the way. The initial unbiased approach taken in this section therefore serves to generate hypotheses for testing by a more rigorous investigation of the protonation state-dependent conformational changes. To this end, we set out to employ enhanced sampling simulations for obtaining conformational free energy landscapes in dependence on a range of protonation state and substrate binding conditions.

### 2D-PMFs show proton-dependent driving forces of PepT2 alternating access

To overcome the time-scale limitations of MD simulations and sample important slow degrees of freedom, many enhanced sampling approaches have been developed (***Hénin et al., 2022***) and employed in the computational study of membrane proteins. (***Harpole and Delemotte, 2018***) An important class of methods that includes (among others) the popular techniques of steered MD (SMD) (***Izrailev et al., 1999***), umbrella sampling (***Torrie and Valleau, 1977***), metadynamics (***Barducci et al., 2008***), adaptive biasing force (ABF) (***Darve et al., 2008***) and the accelerated weight histogram method (AWH) (***Lindahl et al., 2014***) uses a small number of collective variables (CVs) along which to bias the simulation, thus improving exploration of important regions of phase space if the CV includes the relevant slow degrees of freedom (DOFs). If the CV is not optimal, problems can manifest in the form of hysteresis (starting-state dependence) when moving between known end-states. (***Lichtinger and Biggin, 2023***) This is the case for the PepT2 OCC↔OF transition with the simple tip-CV illustrated in figure 2a. Using either metadynamics or SMD with replica-exchange umbrella sampling (REUS), the end-state from which the simulations were started is always favoured in the resulting potential of mean force (PMF), with the hysteresis effect totalling ≈ 40 kcal mol^−1^ for each method (figure S6).

We have recently developed a strategy to overcome such hysteresis issues in conformational sampling which we call MEMENTO (Morphing Endstates by Modelling Ensembles with iNdependent TOpologies, ***Lichtinger and Biggin (2023)***), and have implemented it as the freely available and documented PyMEMENTO package. MEMENTO generates paths between known end-states by coordinate morphing followed by fixing the geometries of un-physical morphed intermediates. Since these paths by definition connect the correct end-states (unlike biased MD methods like SMD, where not reaching the target state in slow orthogonal DOFs is a common source of hysteresis), they can drastically reduce hysteresis in enhanced sampling compared to SMD as a path generation method. We have shown this for several validation cases, including a large-scale conformational change in the bacterial membrane transporter LeuT. After running initial 1D-REUS from MEMENTO replicates along a simple CV guess, we can use the generated MD data to iteratively improve CVs using principal component analysis (PCA), thereby capturing slow motions from long end-state sampling that propagate through MEMENTO as differences between replicates.

Here, we ran triplicates of MEMENTO for the OCC↔OF (standard protonation states and H87&D342 protonated) and OCC↔IF (standard protonation states and E53 protonated) conformational changes, followed initially by 1D-REUS along the tip-CV (figure S7). The results are much more consistent than SMD or metadynamics along the same CV, and the shapes of the PMFs fit well with the trends we previously observed in unbiased MD from the OCC state. Since, however, significant differences between replicates remained, we used principal component analysis (PCA, see Materials and Methods for details) on the sampling collected of the standard protonation state transitions to derive sets of 2-dimensional CVs (figure S8 and supplementary videos 1–4) that capture the main gate-opening motions in the first PC, and the direction along which the differing replicates can be best separated out as the second PC (these correspond to cleft sliding and twisting motions).

Equipped with these CVs, we first studied the protonation-state dependence of the OCC↔OF transition. As figure 3 shows (2D-PMFs in part a, projections onto PC 1 in part b), the OCC state in standard protonation states forms a basin that is metastable with respect to OF (lying ≈ 3 kcal mol^−1^ higher than OF, separated by a barrier of ≈ 3 kcal mol^−1^). Protonation of H87 still leads to a metastable OCC basin, although it is raised by ≈ 4 kcal mol^−1^ and the barrier is decreased to ≈ 1.5 kcal mol^−1^. Protonating D342, in turn, does not affect the relative free energies of the OCC and OF states, but does lower the transition barrier by ≈ 1 kcal mol^−1^. These effects are additive, so that protonation of both H87 and D342 abolishes the metastable OCC state — an observation which agrees with the ability of the OCC state thus protonated to spontaneously transition to OF in unbiased MD (see the Discussion for a comparison with the results obtained by ***Parker et al. (2017)*** on PepT_So_ on this point).

We have also computed 2D-PMFs in further protonation-state and mutation conditions to gain a better understanding of how the H87 and D342–R206 interaction networks control the extracellular gate (figure S9 for the 2D-PMFs, all 1D-reprojections are shown for reference in figure S10). From the data presented thus far, it is not clear whether the effect of H87 pronation on the OCC → OF transition is due merely to the loss of hydrogen-bond interactions with S321, or whether the introduction of a positive charge in this location makes a significant mechanistic contribution. To address this question, instead of protonating H87 we mutated it to alanine (H87A). In the resulting PMF, the OCC state is raised less with respect to OF compared to the protonated version, and the transition barrier increases to ≈ 5 kcal mol^−1^, suggesting that there exists an interaction made by positively charged H87 that becomes particularly relevant in the transition region. Further analysis of the H87 interaction networks in our 2D-REUS trajectories (figure S11a) reveals that when H87 is protonated, the interaction with S321 is substituted by an interaction with D317 that is strongest in the transition region. Interestingly, this suggests an alternative mechanistic role for the essential D317, which — as discussed above — we have not found forming the structurally observed saltbridge with K64 consistently in our simulations.

An equivalent investigation of the D342A mutation results in a PMF that shows both a decrease in the transition barrier, and — as opposed to D342 protonation — also raises the OCC state in energy with respect to OF. This may be explained by the fact that protonated D342 can still hydrogen-bond with R206, so although the interaction is less prominent it presumably still contributes somewhat to OCC state stability (figure S11b). As a control, we also show that although the protonation of the ExxER glutamates (E53 and E56) can affect the relative OCC vs OF free energies, there is no lowering of the transition barrier (while they do have an effect on the transition barrier separating OCC from IF, as discussed below). Convergence analysis and representative 2D-REUS histograms for our OCC ↔ OF PMFs can be found in figures S12 and S13.

We next employed an equivalent approach to investigate the OCC↔IF transition. As can be seen from figure 4, in standard protonation states, the OCC state forms a well-defined basin, connecting to a broader and shallower IF basin raised ≈ 2 kcal mol^−1^ over OCC via a barrier of just over ≈ 3 kcal mol^−1^. When either E53 or E622 are protonated, IF drops to a similar free energy level as OCC and the transition barrier lowers by ≈ 1 kcal mol^−1^. These effects are additive, so that when both E53 and E622 are protonated, the IF state is favoured by ≈ 3.5 kcal mol^−1^, and is accessible from OCC via a barrier of only ≈ 1 kcal mol^−1^. As for the OCC↔OF transitions, these results explain the behaviour we had previously observed in the unbiased MD of figure 2c. The stochastic partial intracellular gate opening seen with those runs can be rationalised through the lower transition barrier from OCC to IF compared to the transition to OF in our PMFs, together with the broad and flat shape of the IF-state basin. Additionally, we have also computed all the equivalents of these PMFs in the presence of Ala-Phe substrate (figure S14), which we will discuss in the section below. All projections onto PC 1 are shown in figure S15, and convergence analysis is provided in figure S16. Taken together, the PepT2 apo 2D-PMFs provide a detailed view of the way the alternating access cycle is driven by proton movement from the extracellular to the intracellular side of the transporter that fits well with its biological function of using two protons from the extracellular medium to energise cycling from OF to IF, and back spontaneously once the protons have left (see the Discussion below).

### Substrate coupling of alternating access includes several distinct mechanisms

Given the evidence presented so far, which provides a plausible model for how protons drive alternating access based just on an investigation of the apo states, it remains unclear how coupling of proton transport to the substrate is achieved — that is, why the transporter cannot be driven by protons without the presence of substrate (and would thus just leak protons). To investigate the mechanism underpinning peptide–proton coupling, we constructed simulation boxes that included a bound Ala-Phe substrate molecule (see Materials and Methods for details). We then calculated the Ala-Phe affinity using absolute binding free energy (ABFE) simulations in different protein states (table 1). Firstly, we observed that the affinity is similar in the OF and IF states, indicating that the binding of substrate alone does not thermodynamically drive the transporter from OF to IF. We did find, however, that protonation of E622 (i.e., the saltbridge parter of the substate N-terminus) significantly decreases substrate affinity. Given that protonation of E622 also favours the OCC→IF transition, we suggest a dual function of E622 protonation that includes both stabilising the IF state with respect to OCC and facilitating substrate release from holo IF.

**Table 1.** Results of ABFE calculations, showing that the affinity of Ala-Phe substrate does not depend much on the conformational state (OF vs IF), but is signicantly decreased on E622 protonation.

| Protein state | AF dipeptide affinity / kcal mol<sup>-1</sup> |
| --- | --- |
| OF | 8.0 ± 0.3 |
| IF | 7.0 ± 0.4 |
| IF & E622 prot | 2.9 ± 0.2 |

To explore what effect the substrate has on the PepT2 conformational landscape, we repeated a set of our 2D-PMFs in the presence of substrate (figure 5). For both the OCC↔OF and OCC↔IF transitions, the 2D-PMFs have similar shapes in the apo and holo states. For OCC↔OF, however, we found an increased width of the OCC state basin in the direction of PC 2, which — as is evident after projection onto PC 1 — stabilises OCC with respect to OF by ≈1 kcal mol^−1^ as an effect of increased OCC flexibility in the orthogonal DOF. We find a similar stabilisation when E53 or E56 are protonated, but not when both H87 and D342 are protonated (figure S9 and S10). This indicates that the presence of substrate in conformations approaching the OCC state from OF may trigger proton movement further down into the transporter — driven by entropic gains from increased flexibility in orthogonal DOFs.

**Figure 5.**
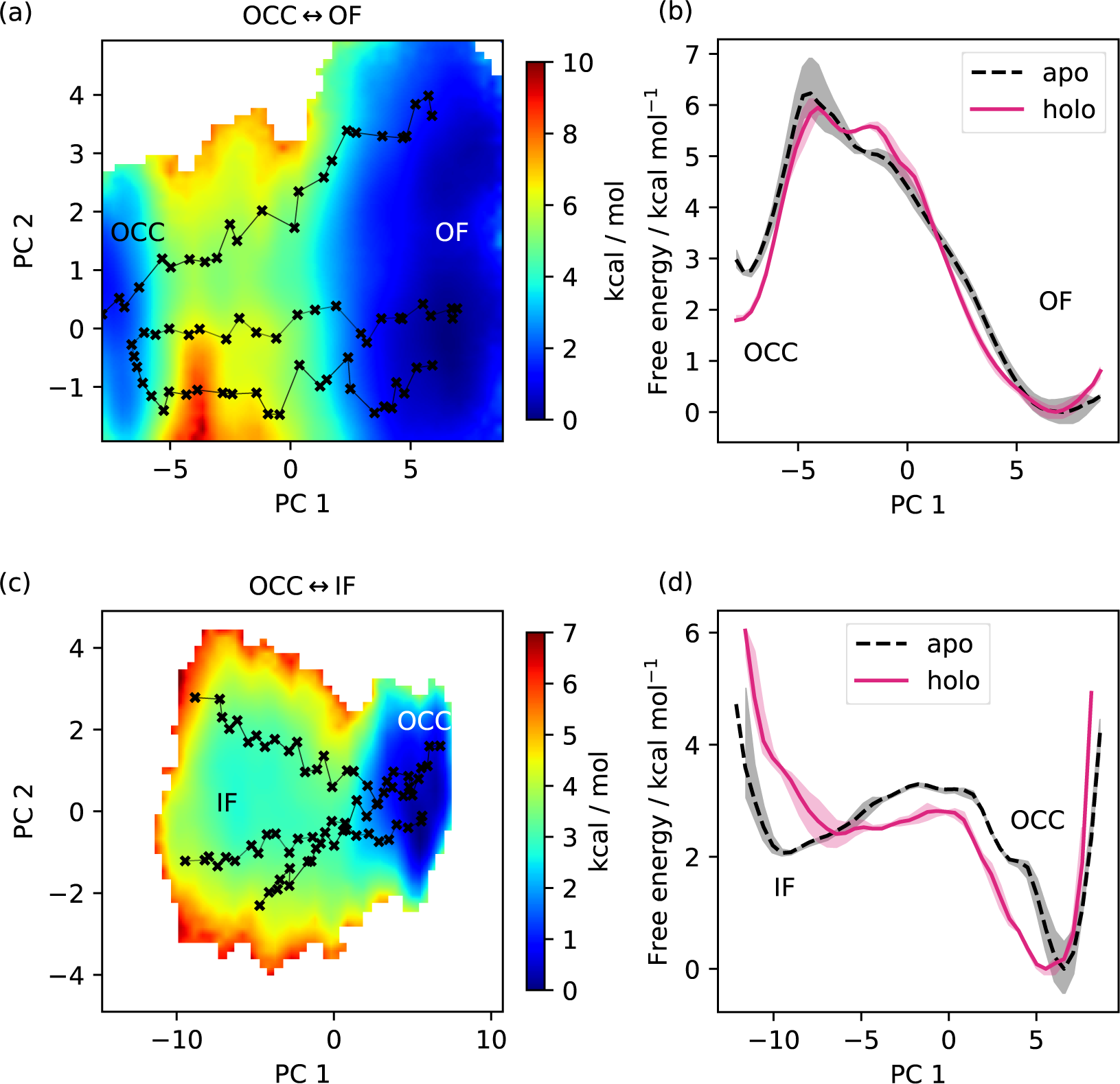
(a) 2D-PMF for the OCC↔OF transition from REUS starting with Ala-Phe-bound PepT2 MEMENTO paths. The OCC state has an increased basin width in PC 2 (compare for figure 3a), and a transition path shifted in PC 2. (b) Projection of the PMF from panel a onto PC 1, showing how in holo PepT2, the OCC state is stabilised by ≈ 1 kcal mol^−1^. Shaded areas indicate convergence errors as the range of PMF values for a given CV value obtained with the first 40 %, the last 40 % and 100 % of sampling included (after alignment to the 100 % curve). Note that the individual PMFs are only determined by our REUS approach up to additive constants, and are shown aligned here at the OF state for convenience of comparison. (c) 2D-PMF for the OCC↔IF transition from REUS starting with Ala-Phe-bound PepT2 MEMENTO paths. The structure of the IF plateau is not significantly affected, but OCC is more flexibile in PC 1. (d) Projection of the PMF from panel c onto PC 1, showing how in holo PepT2, the OCC state has a broader basin, corresponding to intracellular-gate flexibility. Convergence error and alignment of PMFs is shown as in panel b.

The OCC↔IF PMF also presents a broader OCC basin in the presence of substrate, this time in the direction of PC 1 (consistent with the higher OCC-state flexibility directed towards IF observed in unbiased MD, see figure 2c). In the projection onto PC 1, this manifests as a broader free energy well and a lower barrier towards IF, even if the relative energies of the basins are not signicantly affected. This effect is similarly apparent when E53 is protonated, but not with E622 protonation, which instead leads to a raised transition barrier (these opposing trends continue to be additive when both residues are protonated; figure S14 and S15). We reason that this is due to a more flexible substrate orientation (disengaging the N-terminus) when E622 is neutral. While, taking these observations together, substrate binding does lead to an OCC→IF bias, it also seems unlikely that E622 is protonated at the moment when the OCC→IF transition happens in the holo transporter. This may suggest a further intermediate protonation step that our simulations have not captured (see our Discussion below).

As noted above, another possibility for the substrate to engage with the transport cycle is found in R57 interacting both with the substrate C-terminus and with the ExxER glutamates, protonation of which drives the transporter towards the IF state, as our PMFs demonstrate. A natural hypothesis then is that substrate binding — which engages R57 — loosens the R57 interaction with E53 and/or E56, thus enabling the protonation of those residues and progress along the alternating access cycle. To test this hypothesis, we conducted triplicate constant-pH simulations (CpHMD) (***Swails et al., 2014***) to probe the E53 and E56 pKa values in all combinations of OCC / OF and apo / holo conditions (figure 6a). Concerning the E53 pKa value, we see a potential response to substrate binding in the OF state (though the error bars calculated from the triplicate standard deviations overlap) but not in OCC. On the other hand, we do see a raising of the E56 pKa beyond error in holo OF or OCC states compared to apo, amounting to ≈0.6 log units in both cases. As shown in figure S17 and S18, the pKa values estimated for successive data chunks across the CpHMD trajectories vary significantly with simulation time in a complex superposition of timescales, and with a dynamic range larger than the replicate error bars. As an alternative to the replicate-based representation in figure 6, we have therefore also analysed the pooled data for each condition as histograms of pKa values estimated from short chunks of our simulations (figure S19). From this, we recover the same effect of substrate binding on the E56 pKa in the OF and OCC states, as well as a potential effect on the E53 pKa in the OF state only. Since the shift in E56 pKa was more robust across conformational states, we focus on this residue in the following validation and discussion, although we note that if there was in fact a significant raising of the E53 pKa as well, this would further strengthen our conclusions about substrate coupling to ExxER motif protonation.

**Figure 6.**
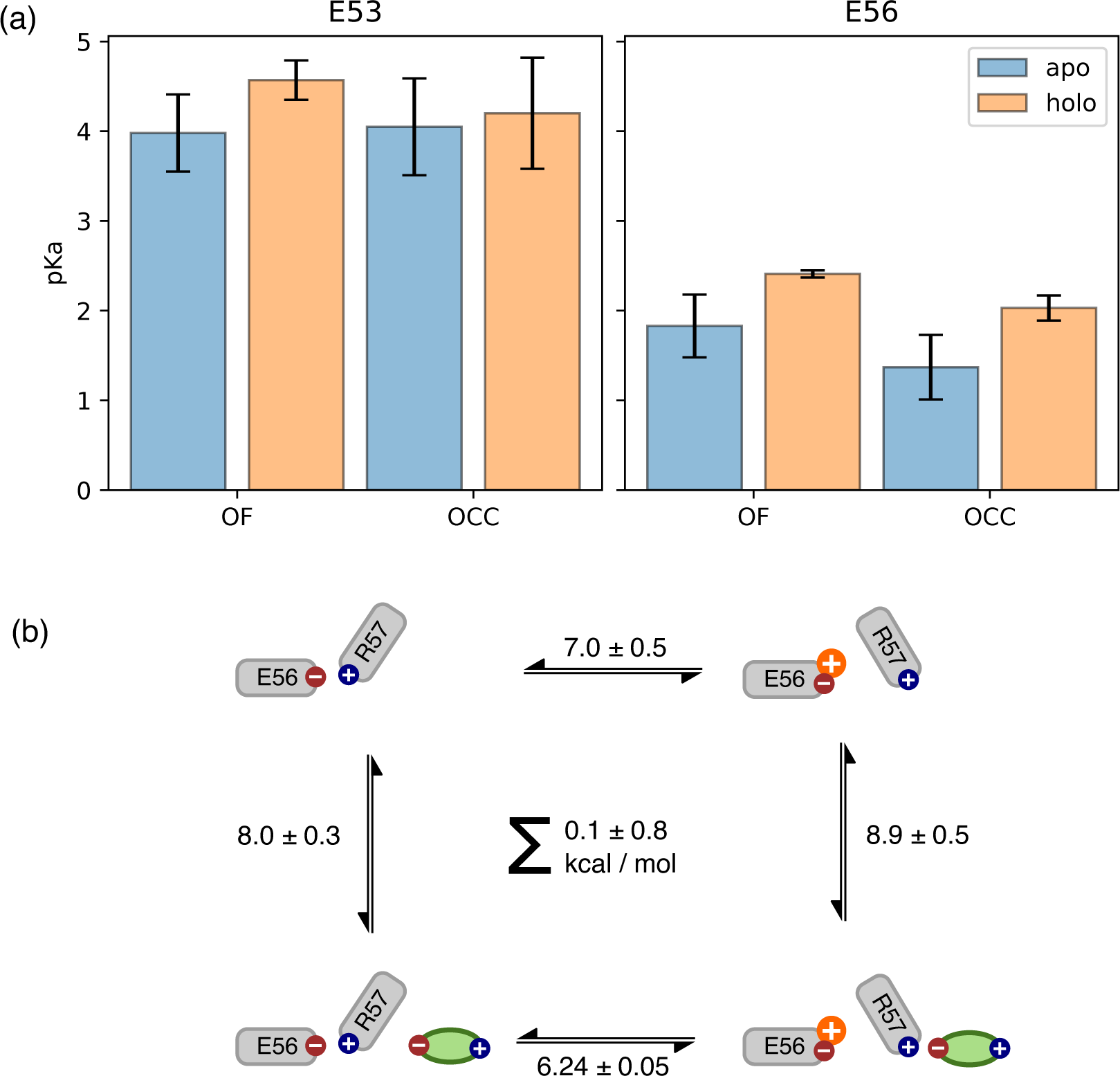
(a) E53 and E56 pKa values from constant-pH MD simulations, in the apo and holo as well as the OF and OCC states, estimated as mean ± standard deviation from triplicate runs (using the full simulation data for fitting the titration curves). The presence of substrate raises the E56 pKa in either conformational state, while some effect on the E53 pKa may also exist in the OF state. (b) Illustration of a themodynamic cycle of E56 protonation and Ala-Phe binding, with edges filled in via CpHMD (converted into kcal mol^−1^ at pH 7) for the top and bottom transitions, and ABFE for the left and right edges. Notably, ABFE displays a response of Ala-Phe affinity to E56 that is consistent with the CpHMD results, and the cycle closes very well. The error in the cycle closure residual is estimated as a square root of the sum of squared standard deviations of the individual edges.

It is important to recognise here that the hybrid-solvent CpHMD method as implemented in AMBER is not rigorous for membrane proteins, since the membrane is not taken into account while evaluating proposed protonation state changes in implicit solvent. On the other hand, we were not able in initial trials to obtain sufficient transition counts to converge an alternative explicit-solvent CpHMD method as implemented in GROMACS. (***Aho et al., 2022***) To validate our results, we therefore also constructed a thermodynamic cycle of substrate binding and E56 protonation in the OF state by including data from separate ABFE calculations as connecting thermodynamic legs (figure 6b). The ABFE results show a complementary effect of the E56 protonation state on substrate affinity, closing the thermodynamic cycle and validating our CpHMD simulations using an orthogonal MD technique. We conclude that substrate binding does indeed facilitate protonation of E56 (whence, we stipulate, the proton moves to E53, which is situated in close proximity). If the neglected membrane environment in the hybrid-solvent CpHMD did produce significant artifacts in the pKa values, then it would appear that there is error cancellation when assessing the impact of substrate binding as a difference of pKa values in the apo and holo conditions.

It should be noted that — as throughout this study, see the discussion below — in studying the coupling between substrate binding and protonation-state changes at the ExxER motif we have not made the substrate C-terminus protonatable. Since, in order to induce E56 protonation, the substrate C-terminus needs to engage R57 in a salt bridge, its pKa is likely to be low, rendering the assumption reasonable for those substrate conformations. However, it is possible that the system could also adopt states in which the ExxER motif salt-bridge network is stable in a way similar to the apo condition while the substrate gets protonated when oriented away from R57. If such conformations contribute significantly to this semi-grand canonical ensemble, the ExxER motif pKa values estimated without taking them into account may exhibit some bias. By undersampling more apo-like conformations in the holo state in this way, it is possible that the calculations presented here overestimate the substrate-induced pKa shift of E56, although the direction of change would be expected to be the same, because the substrate can still engage R57 when it is deprotonated (we speculate that the histograms in figure S19 for the holo state may become bi-modal in this case). While the possibility would need to be taken into account for a more rigorous quantitative estimate of the ExxER pKa value shifts, the pKa calculations would become much harder to converge since the slow degree of freedom of substrate re-orientation would need to equilibrate to the protonation-state changes (which happen fast since they are treated as Monte-Carlo moves in an implicit solvent at regular intervals). Here, we contend ourselves with the more qualitative observation that an appropriately positioned substrate in the canonical, structurally observed binding pose facilitates ExxER-motif protonation.

### Validation in cell-based transport assays

To experimentally validate the results of an MD investigation, a first step is to probe the importance of the key implicated residues for protein function. We note that the literature already contains ample data to show that E53 (***Solcan et al., 2012***; ***Doki et al., 2013***; ***Sun et al., 2014***; ***Jørgensen et al., 2015***; ***Parker et al., 2017***), E56 (***Solcan et al., 2012***; ***Jørgensen et al., 2015***; ***Sun et al., 2014***; ***Zhao et al., 2014***; ***Parker et al., 2017***), R57 (or the equivalent lysine) (***Solcan et al., 2012***; ***Guettou et al., 2013***; ***Doki et al., 2013***; ***Jørgensen et al., 2015***; ***Lyons et al., 2014***; ***Parker and Newstead, 2014***; ***Sun et al., 2014***; ***Parker et al., 2017***; Martinez Molledo et al., 2018), H87 (for those homologues which conserve it) (***Fei et al., 1997***; ***Chen et al., 2000***; ***Newstead et al., 2011***; ***Uchiyama et al., 2003***; ***Parker et al., 2017***), D317 (or the equivalent glutamate) (***Solcan et al., 2012***; ***Doki et al., 2013***; ***Lyons et al., 2014***; ***Parker et al., 2017***; Martinez Molledo et al., 2018; ***Shen et al., 2022***) and E622 (***Solcan et al., 2012***; ***Guettou et al., 2013***; ***Doki et al., 2013***; ***Lyons et al., 2014***; ***Zhao et al., 2014***; ***Parker et al., 2017***; ***Minhas and Newstead, 2019***; ***Shen et al., 2022***) are important for transport through POTs. Of the residues implicated by our simulations, therefore, only S321 and D342 have not been studied before, and thus serve as predictive validation test cases here.

Using cell-based transport assays (see Materials and Methods), we tested the transport activity of rat PepT2 WT and several mutants: H87A (as a positive control known from the literature), I135L (as a negative control, without any expected effect), as well as the mutants of interest D342A and S321A (figure 7, and figure S20 for loading control and membrane localisation micrographs). We note that all our mutants expressed slightly less compared to the WT at the same amount of transfected DNA (0.8 µg), but more than WT at a reduced transfection DNA level (0.5 µg) (figure 7b). To control for this difference in expression levels, we took WT (0.5 µg), which transports ≈20% less than WT (0.8 µg), as a lower bound for the WT transport activity and as the point of comparison for statistical tests. We found that all mutants of residues predicted to be involved in transport displayed significantly reduced transport activity (p-values: 2.2 × 10^−5^ for H87A, 1.6 × 10^−4^ for D342A, 6.9 × 10^−5^ for S321A, while I135L is indistinguishable from WT at p = 0.79). We also note that D342A, although its activity is significantly reduced, still transports more than H87A (p = 3.9 × 10^−7^). This fits well with our 2D-PMF results, where H87 protonation does more than D342 protonation to stabilise OF with respect to OCC.

**Figure 7.**
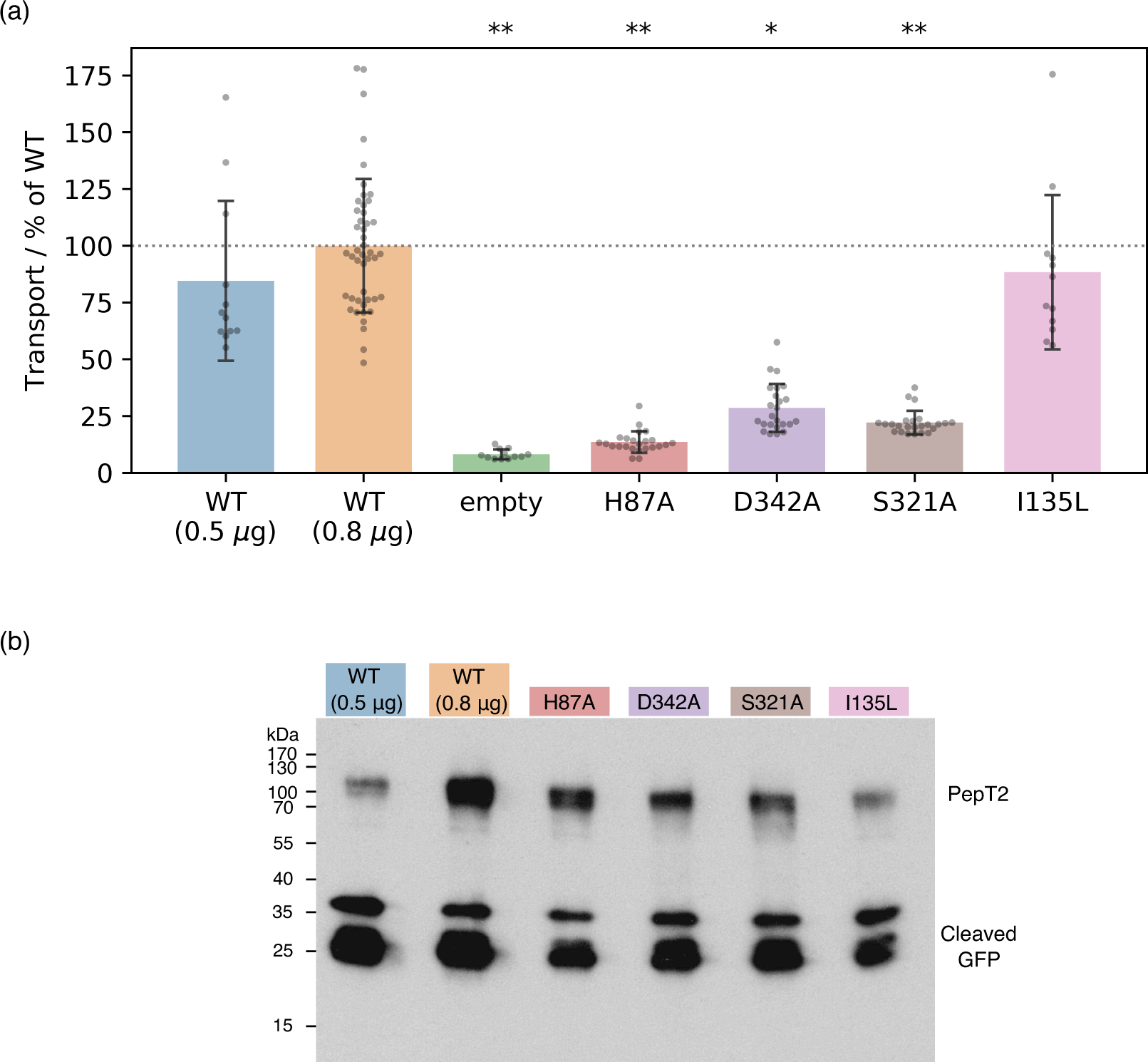
(a) Cell-based transport assays for PepT2 WT (transfected with 0.5 µg, n=12, and 0.8 µg, n=46, of DNA per well), empty plasmid vector (n=12) and PepT2 H87A, D342A, S321A (n=24 each) and I135L (n=12) mutants, all transfected with 0.8 µg of DNA. Diagram shows transport as fluorescence in post-assay lysate divided by total protein concentration, normalised to the WT (0.8 µg) mean. Bars are mean values plus minus standard deviation, and swarm plots samples corresponding to individual wells. Single asterisk indicates *p* < 10^−3^, double asterisks *p* < 10^−4^ significance levels for difference compared to (weaker transporting, 0.5 µg-transfected) WT, as evaluated using a two-tailed t-test. (b) Western-blot showing expression levels of WT and mutant GFP-labelled PepT2, with an anti-GFP primary antibody. All mutants express at levels between the WT transfected with 0.5 µg and 0.8 µg plasmid DNA. Cleaved GFP is also visible at low molecular weight, at levels comparable for WT and mutants.

## Discussion

Integrating the results from extensive sampling across several MD methods, covering all stages of the PepT2 alternating access cycle, we are now in a position to propose a detailed molecular mechanism of the complete transport cycle, including accounts both of proton coupling to conformational changes and of substrate–proton coupling (figure 8). Starting at the apo OCC state without any protons bound, we find in our 2D-PMFs that proton binding to H87 and D342 stabilises OF with respect to OCC (with H87 being the major contributor). Given that H87 and D342 are accessible from the acidic extracellular bulk, and in light of the transition-region stabilisation from the H87 (protonated)–D317 interaction we have identified, our simulations suggest an interpretation where protonation happens in the OCC state, driving the OCC→OF transition by stabilising the OF state over OCC. However, if the OCC→OF transition is kinetically accessible on experimental timescales without prior protonation events (beyond what our MD was able to sample), it would also be consistent with our data that OCC→OF is spontaneous in standard protonation states, and H87 and (possibly) D342 are merely the initial sites of protonation once OF is reached, providing further stabilisation. ^1^ We note in this context the limitation of using non-reactive MD methods for sampling the PepT2 conformational changes. In principle, a multi-dimensional PMF calculated with a reactive MD method where one CV is an explicit proton movement coordinate could disambiguate between the possible scenarios here. However, we believe that such a fully coupled treatment of proton movement and large-scale conformational changes is not yet computationally feasible, and focussed here on achieving convergence of the conformational sampling in discrete protonation-state combinations. The question of whether proton binding happens in OCC or OF therefore warrants further investigation, and indeed the co-existence of several mechanisms may be plausible. Nonetheless, this study contributes important details to a mechanistic understanding of the thermodynamics of proton-coupled alternating access.

**Figure 8.**
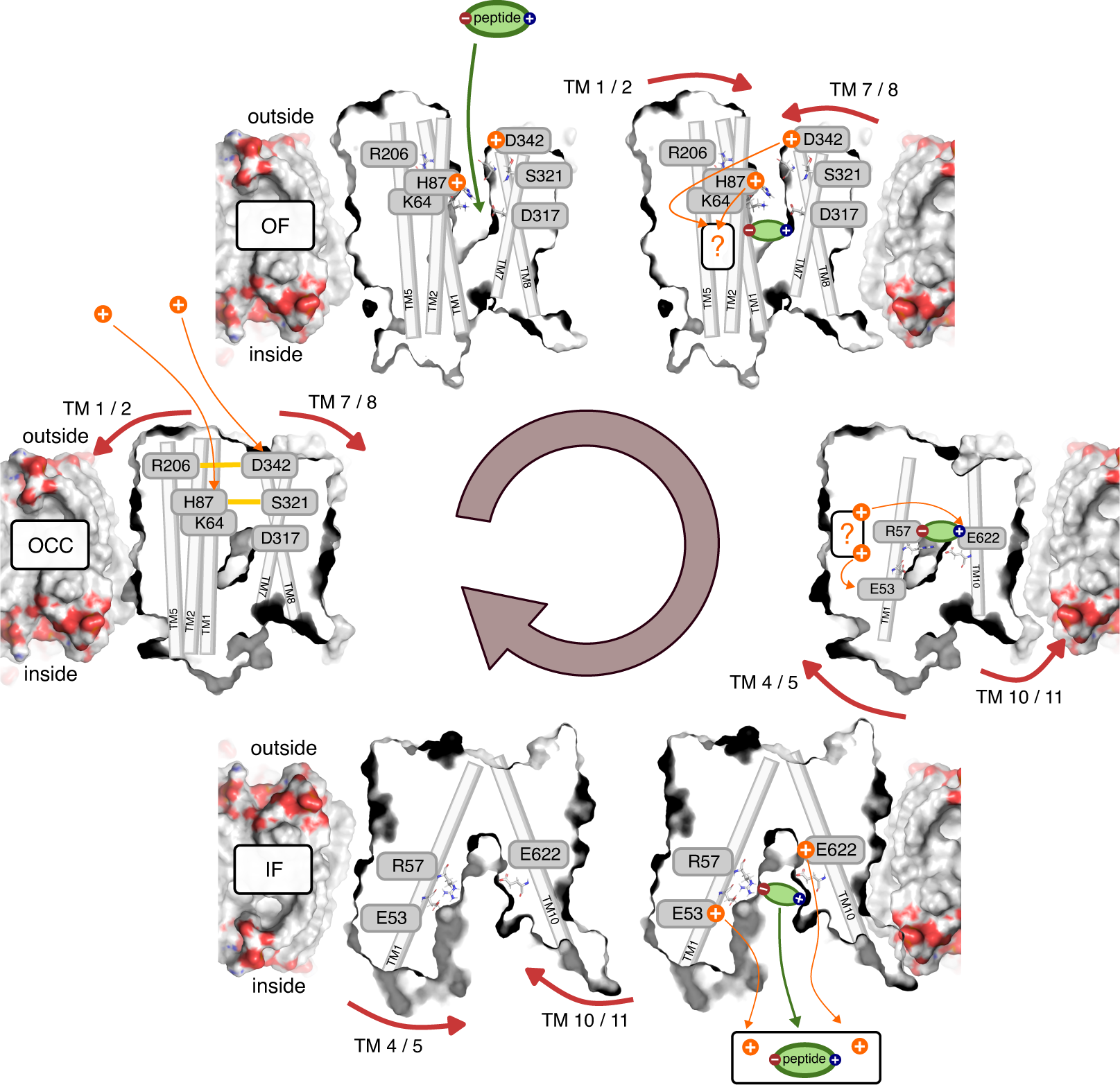
Schematic overview of the PepT2 alternating-access transport cycle proposed in this work. Protons located at a question mark indicate a proton-transfer step with an as-of-yet unknown mechanism regarding intermediate residues.

Our results with regards to the driving forces behind the OCC→OF motion agree with the work of ***Parker et al. (2017)***, who found spontaneous opening of PepT_Xc_ towards OF after protonating the equivalent histidine to H87 (PepT2). In transporters which do not conserve the mammalian histidine at the TM 2 position such as PepT_St_ (***Batista et al., 2019***) and PepT_Sh_ (***Li et al., 2022***), on the other hand, previous simulation studies have implicated protonation of the glutamates equivalent to D317 (PepT2) in the opening of the extracellular gate. Our simulations suggest that the involvement of D317 in the extracellular gating mechanism of PepT2 is by interacting with the protonated H87 to stabilise the transition region of extracellular gate opening. The mechanism of extracellular gating is therefore conserved less widely than the overall alternating access mechanism. This point is further highlighted by our results indicating a role of the D342–R206 saltbridge, which is conserved only among mammalian POTs, but not in PepT_So_ and PepT_Xc_ which do have the TM 2 histidine. This may explain why spontaneous opening in unbiased MD was observed by ***Parker et al. (2017)*** for PepT_Xc_ following just H87 protonation, while for PepT2 in this work, protonation of D342 is also required.

Once the transporter is in the OF conformation, the substrate enters the binding site. We have treated this step al-chemically in our ABFE simulations, so that the data presented here is agnostic of the orientation of the entering substrate and the sequence of engagement of the binding-site residues; previous MD simulations have suggested, however, that positioning of the peptide N-terminus precedes the C-terminus moving into place. (***Parker et al., 2021***) Substrate binding then has two distinct effects: firstly, it exerts a small thermodynamic bias towards the OCC state via increased flexibility in degrees of freedom orthogonal to the overall OF→OCC transition. Secondly, through engaging R57, substrate entry increases the pKa value of E56 in the ExxER motif, thus thermodynamically facilitating the movement of protons further down the transporter cleft. We can also speculate that — in addition to this thermodynamic favouring of E56 protonation — there might be a kinetic effect on proton transfer from moving the positively charged R57 out of the way of the incoming proton. This could be investigated using reactive MD or QM/MM simulations (both approaches have been employed for other protonation steps of procaryotic peptide transporters, see ***Parker et al. (2017)*** and Li et al. (2022)), however the putative path is very long (≈ 1.7 nm between H87 and E56) and may or may not involve a large number of intermediate protonatable residues, in addition to binding site water. While such an investigation is possible in principle, it is beyond the scope of the present study. Likewise, a coupled enhanced-sampling treatment involving both proton movement and large-scale conformational changes — as discussed above for the ordering of steps in the OCC → OF transition — would make for interesting future work, once it becomes computationally tractable.

Our data is not fully determinate with respect to whether protons move to E56 before or after the OF→OCC conformational transition (our CpHMD, for example, remains agnostic on this matter since the shift in the E56 pKa value induced by the substrate is evident in both the OCC and OF states). We may interpret the fact that OCC is raised in energy while H87 is protonated and substrate-induced OCC stabilisation is not found when H87 and D342 are protonated (but does occur when E56 is protonated) as an indication that proton movement is favoured before the transition into OCC is complete. On the other hand, the transition-region interaction between protonated H87 and D317 could also be interpreted as a potential facilitator of the OF→OCC transition. We thus speculate that the proton movement processes may happen as an ensemble of different mechanisms, and potentially occur contemporaneously with the conformational change. This, in addition to a flexible binding pocket, may also contribute to the substrate promiscuity mechanism.

We note at this stage that — throughout our study — we have not investigated the possibility of the substrate C-terminus itself becoming (transiently) protonated. This would need to be taken into account when treating proton movement through the transporter explicitly in the future (see the discussion of such approaches above). There is evidence in our simulations that an additional protonation site — aside from H87, D342, E53, E56 and E622 — may be involved in the mechanism, since E622 protonation, while biasing the transporter towards IF, also increases the OCC→IF transition barrier if Ala-Phe substrate is bound (we therefore indicate the proton movements at these stages with a question mark in figure 8). There is thus the intriguing possibility that the substrate itself may temporarily hold the proton, although given the nature of the data presented here only speculation is currently possible on this point. What is clear from our 2D-PMFs, regardless, is that protonation of ExxER does act as an intracellular gate trigger (and may also pull the transporter through the chemical equilibria all the way from OF). Taking together our 2D-PMFs and ABFE simulations, it is also clear that E622, in addition to being essential for peptide recognition, plays two further roles: its protonation both facilitates substrate release and makes an additive contribution to the IF-directed bias exerted by the intracellular gate triggers (whether E622 forms ‘part’ of the intracellular gate remains then as a merely linguistic question). At this stage, we do not yet have an understanding of how exactly intracellular gate opening (which involves breaking an assembly of several hydrophobic residues) is effected by the protonation of these glutamate residues. This question should prove interesting to study in future work. Once the substrate (driven by E622 protonation) and the protons (driven by their electrochemical potential gradient) have left through the open intracellular gate into the intracellular bulk, the resulting apo, standard-protonation-state IF conformation has a thermodynamic preference to return to OCC as evidenced in our 2D-PMFs. We thus arrive back at the starting state and have completed the proton- and substrate-coupled alternating access cycle.

In support of our MD data, we present mutational analysis in cell-based transport assays. Mutations of H87, S321 and D342 to alanine all significantly decrease transport activity, with H87A having the strongest effect. Taken together with similar results in the literature on E53, E56, R57, D317 and E622 (as referenced above), all residues implicated by our study have therefore been confirmed in their importance for transport via mutagenesis. While the cell-based assay used here cannot differentiate for example between proton-coupling and non-proton-coupling residues, our results still provide a useful first step towards the validation of the gating mechanisms we propose with our PMFs, and should prove informative for the future design of more in-depth experiments.

In conclusion, this study utilises the recent wealth of bacterial and mammalian peptide transporter structures to construct a model of their alternating access mechanism. We explain how the movement of two protons through the transporter drives the accompanying conformational changes, as well as how conformational changes and proton-movement events are coupled to the presence of substrate. Questions regarding some of the finer details, notably the precise sequence of proton movements and conformational transitions (if a single such sequence exists) and whether a further protonation site contributes to the mechanism remain open for future investigation. Nonetheless, the evidence supplied here addresses the alternating access proton-symport mechanism in unprecedented detail, particularly through the extensive use of free-energy simulation techniques. This information will prove useful for the project of employing peptide transporters as vehicles for drug delivery — especially since what determines the efficacy of a transporter substrate is not only related to affinity but crucially also to an ability of the substrate to move through the steps of the alternating access cycle once bound to the transporter.

## Methods and Materials

### Data availability

Simulation and experimental data produced in this work is available at doi: 10.5281/zenodo.10561418. This includes key coordinate files and python scripts, as well as simulation trajectories projected onto the CVs of interest in plumed output format, and relevant processed files for PMF, pKa and ABFE calculations. Full trajectories will be shared upon reasonable request. The PyMEMENTO software is freely available at https://github.com/simonlichtinger/PyMEMENTO.

### Definition of tip and base bundle CVs

For the analysis and interpretation of our unbiased MD runs, as well as for the use as a CV for initial metadynamics and umbrella sampling trials, we constructed the tip-CV and base-CV as centre-of-mass distances between the Cα atoms of the top and bottom 11 residues of the N-terminal and C-terminus bundles respectively, as illustrated in figure 2a. These residues were picked as the consensus of the DSSP analysis (***Kabsch and Sander, 1983***) of the PepT2 conformational states derived below, listed in table 2.

**Table 2.** Residue numbers used in the defintion of the tip-CV and base-CV.

| Bundle | Residue numbers |
| --- | --- |
| N-terminal bundle tips | 63–74, 79–90, 120–131, 143–154, 194–205, 217–228 |
| N-terminal bundle bases | 46–57, 93–104, 110–121, 161–172, 177–188, 227–238 |
| C-terminal bundle tips | 320–331, 341–352, 392–403, 609–620, 655–666, 671–682 |
| C-terminal bundle bases | 290–301, 359–370, 376–387, 626–637, 642–653, 686–697 |

### MD setup and equilibration

We obtained protein coordinates from cryo-EM for the OF (7NQK) (***Parker et al., 2021***) and IF-partially-occluded (7PMY) (***Killer et al., 2021***) PepT2 conformations, as well as from alphafold 2 (***Jumper et al., 2021***) for the fully-open IF state. We used MODELLER (***Šali and Blundell, 1993***) to fit the rat PepT2 sequence (as used by ***Parker et al. (2021)***) to the human PepT2 7PMY and alphafold models, using residues 43–409 (TM 1–9) linked as a continuous chain to residues 604–700 (TM 10–12), thereby truncating the extracellular domain as done by ***Parker et al. (2021)*** in their MD simulations. We scored 200 models with QMEANDisCo (***Studer et al., 2020***) and selected the highest-scoring protein model for embedding into a 3:1 POPE:POPG bilayer of target size 10 * 10 nm (210/72 lipid molecules for IF and OCC, 218/72 for OF) with the CHARMM-GUI membrane builder. (***Wu et al., 2014***) We added ACE/NME capping residues using pymol and solvated the membrane system using GROMACS (***Abraham et al., 2015***) with approximately 21,000 solvent molecules (precise number varies between replicates) at a NaCl concentration of 0.15 M in an orthorhombic box of around 9.9 * 9.9 * 10.8 nm side lengths. Topologies were generated using the AMBER ff14.SB (***Maier et al., 2015***) and slipids (***Jämbeck and Lyubartsev, 2012***) forcefields.

Using the GROMACS MD engine (***Abraham et al., 2015***) in versions 2021.3/2021.4 (the slight version discrepancy is because of different installations on two compute clusters we used), we energy-minimized the systems, assigned initial velocities, and equilibrated with Cα-atom restraints for 200 ps in the NVT ensemble with a leap-frog integrator (using the modified v-rescale thermostat with a stochastic term (***Bussi et al., 2007***) at 310 K throughout our work), then 1 ns in the NPT ensemble (with the berendsen barostat), followed by 20 ns of further Cα-restrained NPT equilibration (using the Parrinello-Rahman barostat (***Parrinello and Rahman, 1981***), as for all subsequent production runs).

We obtained Ala-Phe dipeptide-bound boxes by aligning the holo PepT1 cryo-EM structure (7PMW) onto our equilibrated PepT2 MD boxes, copying the ligand coordinates, and repeating the same equilibration protocol as before (where the peptide substrate Cα atoms were also restrained). The peptide ligand was parametrised using AMBER ff14.SB. (***Maier et al., 2015***)

### Derivation of OF, OCC and IF conformational states

As shown in supplementary figure S2a, the IF-partially-occluded structure (7PMY) does not behave well in MD (1 µs production runs from triplicate embeddings), since it either partially opens its extracellular gate (replicates 2–3) or partial helical unfolding in the intracellular gate occurs due to hydrophobic collapse (rep 1). This may be due to a variety of factors; one possibility is instability in the protein following the removal of the bound substrate in our simulations. In contrast (figure S2b), embedding replicates 1 and 3 of the alphafold IF state behave well. We picked the end-coordinates of replicate 3 as our IF state, due to the wider opening of its intracellular gate. We then sought to derive an OCC state via MD from replicate 1, see the paragraph below. We also note that embeddings from the OF cryo-EM structure (7NQK) remain stable in the OF conformation, we picked replicate 1 for our work.

To derive an OCC state from an IF box, we conducted 5 replicates of well-tempered metadynamics (***Barducci et al., 2008***) as implemented in PLUMED 2.7 (***Tribello et al., 2014***) along the base-bundle CV (see figure 2a), using 8 walkers, hill height 1 kJ mol^−1^ with sigma 0.022 nm deposited every 500 steps, and a bias factor of 100. From the resulting set of trajectories, we picked frames around the mark of 20 ns simulation time and a base-CV value of around 2.0 nm (we chose these values based on visual inspection of the trajectories, where we noticed that base-CV values significantly below 2.0 nm lead to artefacts such as partial unfolding of the ends of helices, as did continuing the metadynamics simulations for longer than necessary to obtain the desired states). We ran triplicate 100 ns-long unbiased MD from the obtained states for each of the five replicates, and found the OCC state obtained from the first replicate to be stably situated within the range of base-CV values observed in the OF-state trajectories. The micro-second-long unbiased MD runs as well as our 2D-PMFs confirm that this protein conformation is a stable basin, and that different protonation states of key residues can drive its opening to either the IF or OF states. While this does not rule out the existence of different OCC states, it confirms the properties of the conformation we found as a functional OCC state.

### Unbiased MD of the OCC state

Unbiased MD runs of the OCC state in different conditions were conducted in triplicates using the same simulation parameters as for the long Cα-restrained equilibrations described in the section on equilibration, but removing all restraints. The starting coordinates were — for the first replicate — the OCC state derived as described in the foregoing section, and the second and third replicates were initialised from the 500 ns and the final frame of the first replicate trajectory. Protonation states changes and mutations were carried out using PyMOL and GROMACS pdb2gmx independently for each replicate, followed by re-running the full equilibration protocol for all new boxes. Taken together, we conducted unbiased MD for 24 conditions, giving 72 µs of production sampling.

### Metadynamics and steered MD

For our initial trials of enhanced sampling on PepT2 conformational changes — which showed hysteretic behaviour (see figure S6), we attempted steered MD (SMD) and metadynamics for the OCC↔OF transition in the WT, unprotonated state.

Two instances of 8-walker well-tempered metadynamics were run, starting from the OF and OCC states, biased along the tip-CV with hills of height 1 kJ mol^−1^ and sigma 0.0455 nm deposited every 500 steps, using a bias factor of 100. A harmonic flat-bottom restraint with boundaries of 2.0–3.0 nm and force constant 5 × 10^4^ kJ mol^−1^ nm^−2^ was applied on the CV value. Sampling was run for 108 ns per walker for the simulations starting from OF, and 213 ns per walker starting from OCC.

To generate paths for umbrella sampling, SMD was run starting at OF towards OCC and vice versa with the heavy-atom RMSD to the target conformation as CV, using a harmonic potential centered to zero RMSD with a force constant sliding from 0 up to 2.5 × 10^5^ kJ mol^−1^ nm^−2^ over 200 ns. The harmonic potential was then switched off over 2 ns, followed by 48 ns of unbiased MD. We then projected the SMD trajectories onto the tip-CV, picked 48 frames spaced equally along the CV and performed 1D replica-exchange umbrella sampling (REUS) using a force constant of 3 × 10^4^ kJ mol^−1^ nm^−2^, for 92 ns per window for the OCC→OF derived boxes and 127 ns per window in the reverse direction. A total of 13.6 µs of MD time was thus expended on these trials.

### MEMENTO path generation

We have recently proposed the MEMENTO method for history-independent path generation between given end-states. (***Lichtinger and Biggin, 2023***) In short, protein coordinates are morphed, followed by reconstructing an ensemble of structures at each morphing intermediate using MODELLER. Monte-Carlo simulated annealing with an energy function based on between-intermediate RMSD values then finds a smooth path through these ensembles. For membrane proteins, lipids are taken from the end-state that occupies a larger area in the membrane (in this case, OF for the OCC↔OF transition, and IF for OCC↔IF), and fitted around the new protein coordinates by expanding the membrane, followed by iterative compression and energy minimization. Ligands are not morphed but translated, interpolating between the ligand centres of masses in the end-states, and then equilibrated in the protein structure using MD. MEMENTO is implemented as the PyMEMENTO package (https://github.com/simonlichtinger/PyMEMENTO), we provide example scripts for its usage on PepT2 in the supplementary data.

In this study, we ran MEMENTO with 24 windows in triplicates for both the OCC↔OF and OCC↔IF transitions in different protein protonation and mutation states, and in the presence or absence of ligands. The apo and holo MEMENTO replicates were initialised from the 0 ns, 500 ns and 1000 ns frames of the first replicate of the 1 µs unbiased MD run for each conformational state (using always the unprotonated, WT condition, but apo/holo trajectories respectively). Protonation state changes and mutations were then carried out using the built-in functionality of PyMEMENTO, and equilibrated at each intermediate state for 90 ns. The total MD simulation time spent on equilibrations as part of the MEMENTO method across the 22 sampled conditions was 47 µs.

### 1D-PMF calculations

Starting with the triplicate equilibrated MEMENTO boxes for the (all apo) OCC↔OF standard protonation and H87 & D342 protonated states, as well as OCC↔IF standard protonation and E53 protonated states, we ran 1D-replica exchange umbrella sampling (REUS, exchange every 1000 steps; using PLUMED 2, ***Tribello et al. (2014)***) along the tip- and base-CVs respectively, using a force constant of 4 × 10^3^ kJ mol^−1^ nm^−2^. The amount of sampling collected in each case is summarised in table 3.

**Table 3.**
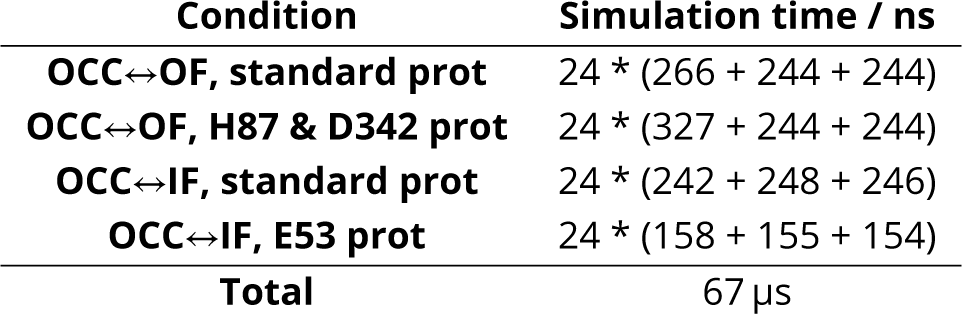
Overview of all 1D-PMF sampling.

### 2D-CV derivation

Using the trajectory data from our 1D-PMFs, we derived 2D CVs via a PCA-based approach we have previously described for LEUT. (***Lichtinger and Biggin, 2023***) We pooled all sampling collected in 1D-REUS runs along the tip-CV for apo OCC↔OF (and equivalently for OCC↔IF. The same procedure was taken for these trajectories, and we will only explicitly write about OCC↔OF in the following paragraph). Using GROMACS tools, we ran PCA of the Cα positions of residues contained in the transmembrane region (see table 2 above). The first princial component (PC) accounts for 50% of the variance; adding an extra 15 PCs increases coverage to 78% of the variance (comparable to our results on LEUT). The first PC (see figure S8) describes the gating motions of the respective conformational changes, behaving similarly to the tip CV (or base CV) — expectedly so, given it was the CV used in our 1D-REUS. To explain differences between replicates (see figure S7a and c), we used differential evolution (***Storn and Price, 1997***) as implemented in scipy (***Virtanen et al., 2020***) to maximise an entropy-like metric of distances between MEMENTO path frames for linear combinations of the PCs 2–16:

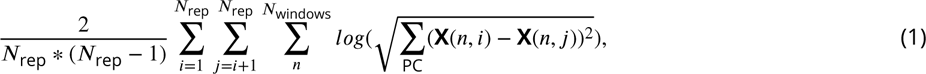

where *N*_rep_ is the number of replicates and by X(*n*, *i*) − X(*n*, *j*) we denote the distance between two conformational frames in different replicates *i* and *j*, evaluated in a projection along a given combination of principal components. The result was termed PC 2 henceforth for simplicity and used as the second CV in 2D-REUS (figure S8).

### 2D-PMF calculations

Using the same equilibrated MEMENTO paths as above and the 2D-CVs we derived, we calculated 2D-PMFs of the OCC↔OF and OCC↔IF transitions in several protonation / mutation states, with and without Ala-Phe substrate bound, using 2D-REUS. As shown in figure S12, we found that a lower force constant of 2 × 10^6^ kJ mol^−1^ nm^−2^ leads to good histogram overlap in the lower-lying regions of the PMF, but has poor overlap near the transition region. In turn, a higher force constant of 1 × 10^7^ kJ mol^−1^ nm^−2^ gives good window overlap in the transition region while not sampling broadly enough in large basins. Therefore, for each condition and MEMENTO replicate, we ran windows at both force constants and included them all in the WHAM analysis, thus ensuring sufficient sampling through the CV space. Replica-exchange was run within the 24 windows corresponding to each MEMENTO replicate–force constant combination, and a total of 144 windows contribute to each 2D-PMF.

The sampling collected in all conditions is detailed in table 4, aiming for between 180 and 210 ns per window though exact amounts differ with heterogeneous hardware and slightly different box sizes.

**Table 4.** Overview of all 2D-PMF sampling.

| Condition | Simulation time / ns |
| --- | --- |
| OCC↔OF, standard prot | 24 * (233 + 195 + 195 + 208 + 208 + 207) |
| OCC↔OF, H87 prot | 24 * (230 + 200 + 188 + 201 + 228 + 229) |
| OCC↔OF, D342 prot | 24 * (204 + 232 + 204 + 224 + 229 + 207) |
| OCC↔OF, H87&D342 prot | 24 * (196 + 196 + 195 + 194 + 194 + 201) |
| OCC↔OF, D342A | 24 * (182 + 183 + 206 + 181 + 183 + 183) |
| OCC↔OF, H87A | 24 * (176 + 182 + 183 + 184 + 182 + 177) |
| OCC↔OF, E53 prot | 24 * (185 + 180 + 209 + 179 + 218 + 213) |
| OCC↔OF, E56 prot | 24 * (194 + 194 + 194 + 192 + 193 + 193) |
| OCC↔OF, R206D&D342R | 24 * (227 + 213 + 183 + 222 + 211 + 180) |
| OCC↔OF, holo, standard prot | 24 * (200 + 196 + 195 + 200 + 194 + 201) |
| OCC↔OF, holo, E53 prot | 24 * (185 + 180 + 210 + 179 + 218 + 213) |
| OCC↔OF, holo, E56 prot | 24 * (194 + 194 + 194 + 192 + 193 + 193) |
| OCC↔OF, holo, H87&D342 prot | 24 * (198 + 191 + 198 + 190 + 182 + 201) |
| OCC↔IF, standard prot | 24 * (197 + 216 + 203 + 192 + 197 + 199) |
| OCC↔IF, E53 prot | 24 * (197 + 208 + 199 + 192 + 193 + 195) |
| OCC↔IF, E622 prot | 24 * (199 + 201 + 200 + 196 + 217 + 197) |
| OCC↔IF, E53&E622 prot | 24 * (193 + 237 + 240 + 187 + 250 + 200) |
| OCC↔IF, holo, standard prot | 24 * (190 + 195 + 191 + 196 + 192 + 197) |
| OCC↔IF, holo, E53 prot | 24 * (189 + 196 + 197 + 199 + 189 + 198) |
| OCC↔IF, holo, E622 prot | 24 * (194 + 191 + 206 + 197 + 198 + 199) |
| OCC↔IF, holo, E53&E622 prot | 24 * (197 + 198 + 180 + 195 + 190 + 204) |
| Total | 598 μs |

### Absolute binding free energies

To probe the affinity of the Ala-Phe substrate to the PepT2 OF and IF conformations in different protonation states, we conducted absolute binding free energy (ABFE) simulations (***Aldeghi et al., 2016***, ***2018***) in gromacs, using an equilibrium approach. For this, we changed our mdp files to use the stochastic dynamics integrator (doubling as a thermostat) and set the relevant free-energy flags, including soft-core van-der-Waals interactions (alpha = 0.5, power = 1, sigma = 0.3) and the couple-intramol=yes flag for consistency with larger ligands in other work. Our lambda-protocol was to first add Boresch restraints (***Boresch et al., 2003***) (for the complex thermodynamic leg only, ligand side was calculated using the analytic formula; through values 0, 0.01, 0.025, 0.05, 0.075, 0.1, 0.2, 0.3, 0.4, 0.5, 0.6, 0.8, 1.0), then annihilate coulumb interactions (even 0.1 spacings) followed by vdw interactions (even 0.05 spacings). We equilibrated for 200 ps of NVT and 1 ns of NPT at each lambda window, and ran production simulations with replica-exchange attempts every 1000 steps for 30 ns per window on the complex thermodynamic leg and 100 ns on the ligand-only leg.

The Boresch restraints for ABFE simulations were obtained MDRestraintsGenerator (***Alibay et al., 2022***) by running the restraint finding algorithm over the ≈200 ns 2D-REUS trajectory at the relevant conformation and protonation state, and we used the trajectory frame closest to the restraint centre as input for subsequent ABFE. This was done for each of the triplicate MEMENTO runs, giving 3 candidate ligand binding poses. For the pose that was found to have the highest single-replicate affinity, four replicates of unbiased, restraint-free 200 ns-long equilibrations were also started from the frames, processed with MDRestraintsGenerator and used to make replicates for ABFE runs to give an error estimate as mean plus-minus standard deviation. By following this protocol for the OF, OF E56 prot, IF and IF E622 prot conditions, we sampled for a total of 4 conditions * 7 boxes per condition * 44 windows * 31.2 ns = 38 µs.

### Constant-pH MD

To probe the substrate-dependence of the E53 and E56 pKa values, we ran constant pH simulations (CpHMD) using the hybrid solvent approach with discrete protonation states (***Swails et al., 2014***) as implemented in the AMBER software. (D.A. ***Case et al., 2021***) We took the MEMENTO starting frames from above as triplicate initial coordinates of OF and OCC states in the presence and absence of Ala-Phe. We then used tleap and in-house scripts to convert our boxes to the AMBER constant pH forcefield fork for protein, substrate and solutes and to the lipid21 force field (***Dickson et al., 2022***) for the membrane. We prepared constant pH simulations as in the tutorial by ***Swails (2014)***, and ran them for 1 µs at 8 pH replica windows (ph 0–7), in the NVT ensemble with a langevin thermostat (at 310 K as before), attempting protonation state changes every 100 steps, running 100 steps of relaxation dynamics for every exchange, and attempting replica exchange every 1000 steps. Analysis was performed using the cphstats programme and in-house scripts for fitting titration curves. In figure 6, analysis is performed per replicate, reporting mean ± standard deviation for each condition and residue. In figure S17–18 we show pKa values estimated over simulation time from 10 ns chunks of all CpHMD runs. We also analyse this data in terms of histograms of the chunk-estimated pKa values, pooling all data for each condition and residue. A total of 4 conditions * 3 replicates * 8 windows * 1 µs = 96 µs of sampling were thus collected.

### Cell-based transport assays

Transport assays were carried out using a modified version of the protocol by ***Parker et al. (2021)***. Hela cells were cultured in DMEM + GlutaMAX medium, supplemented with 10% FBS. 12-well plates were prepared by seeding 9 × 10^4^ cells per well in 1 mL of medium, and transfected after 24h with 0.8 µg of PepT2-constructs in pEF5-FRT-eGFP vector (or 0.5 µg of insert vector + 0.3 µg of empty vector, where specified), with 1.6 µg of fugene transfection reagent. The medium was exchanged 24h post transfection, and assays were carried out 40h post transfection. Cells were washed 2 times with ≈0.6 mL of assay buffer (20 mM HEPES pH 7.5, 120 mM NaCl, 2 mM MgSO_4_ and 25 mM glucose), then incubated with 0.3 mL assay buffer containing 20 mM β-ala-lys-AMCA substrate for 15 min. The cells were then washed 3 times with ≈0.6 mL of assay buffer, and incubated with 0.25 mL of lysis buffer (20 mM Trist pH 7.5 + 0.2% Triton x-100) for 5 min. The fluorescence (340 nm excitation, 460 nm read-out) of 0.15 mL of the lysate was normalised by the protein amount in each well (as determined from BCA assay of 20 µL lysate). We removed two outliers from the WT (0.8 µg) transport assay dataset, giving n=46. The data was then scaled to the mean WT (0.8 µg) transport level as 100%.

### Protein expression controls

For comparing PepT2 WT and mutant expression levels, Hela cells were seeded in 6-well plates at 1.8 × 10^5^ cells per well in 2 mL of medium. Transfection was after 24h with 1.6 µg of PepT2-constructs in pEF5-FRT-eGFP vector (or 1.0 µg of insert vector + 0.6 µg of empty vector, where specified) and 3.2 µg of fugene; the medium was exchanged 24h post transfection. The cells were washed 3 times with ≈0.6 mL of PBS, harvested using 0.1% trypsin, pelleted, re-suspended in 100 µL PBS with protease inhibitor and lysed through 3x freeze-thawing. The lysates from 3 wells were pooled for each mutant to increase between-sample consistency. 4.5 µL of each sample were loaded onto a 10% SDS-PAGE gel, and western blot was performed using an anti-GFP antibody. The membrane was then stripped and developed again with an anti-β-actin antibody to control for gel loading.

### Protein localisation controls

To confirm the plasma membrane localisation of PepT2 WT and mutants, we seeded 1.8 × 10^5^ Hela cells per well in 2 mL of medium in a 6-well plate with added coverslips. Transfection was as for the expression controls 24h after seeding. 20h post transfection, the cells were washed 3 times with ≈0.6 mL of PBS, fixed with PFA for 10 min at room temperature, washed 3 times, incubated with PBS + 50 mM NH_4_Cl for 10 min, washed 3 times, incubated with PBS + 0.1 % Triton x-100 for 5 min, washed 3 times, stained with DAPI, washed 5 times, and mounted on slides with ImmuMount. Images were recorded in the GFP and DAPI channels.

## Supporting information

Video 1 - First Principal Component (PC)of OCC to OF state

Video 2- Second Principal Component (PC)of OCC to OF state

Video 3 - First Principal Component (PC)of OCC to IF state

Video 2 - Second Principal Component (PC)of OCC to IF state

## Acknowledgments

We would like to thank Dr. Zhiyi Wu for general training in molecular dynamics methodology and help in the early stages of this project, Dr. Irfan Alibay for training in ABFE simulations, Dr. Gabriel Kuteyi for training in mammalian cell culture techniques and Sacha Salphati for training in fluorescent microscopy, as well as all members of the Biggin and Newstead groups for helpful discussions.

This project was funded by the Wellcome Trust (Grant ID: 218514/Z/19/Z). Compute resources were also provided by the EPSRC ARCHER2, Jade 2 and N8 CIR BEDE facilities, granted via the High-End Computing Consortium for Biomolecular Simulation (HECBioSim), supported by EPSRC (EP/X035603/1).

## Footnotes

1 It may seem like the latter model is favoured by the fact that in our PMFs, OCC lies higher than OF, even when neither H87 nor D342 are protonated. We believe, however, that there is a danger of overinterpreting this feature of the PMF. Any combination of effects from forcefields, lipid composition and the population shifts afforded by transmembrane electrochemical gradients could perturb the conformational equilibria. It would therefore not be meaningful to interpret the shape of a single PMF in this way; only responses of the PMF to protonation-state or substrate-binding changes should be used to inform our view of the conformational cycle, since these are likely to benefit from error cancellation with respect to factors that act on the overall protein conformations (which are conserved between the different conditions in which the PMFs were sampled).

