## Supplemental Information for "The mechanism of mammalian proton-coupled peptide transporters"

### Supplemetary figures to: The mechanism of mammalian proton-coupled peptide transporters

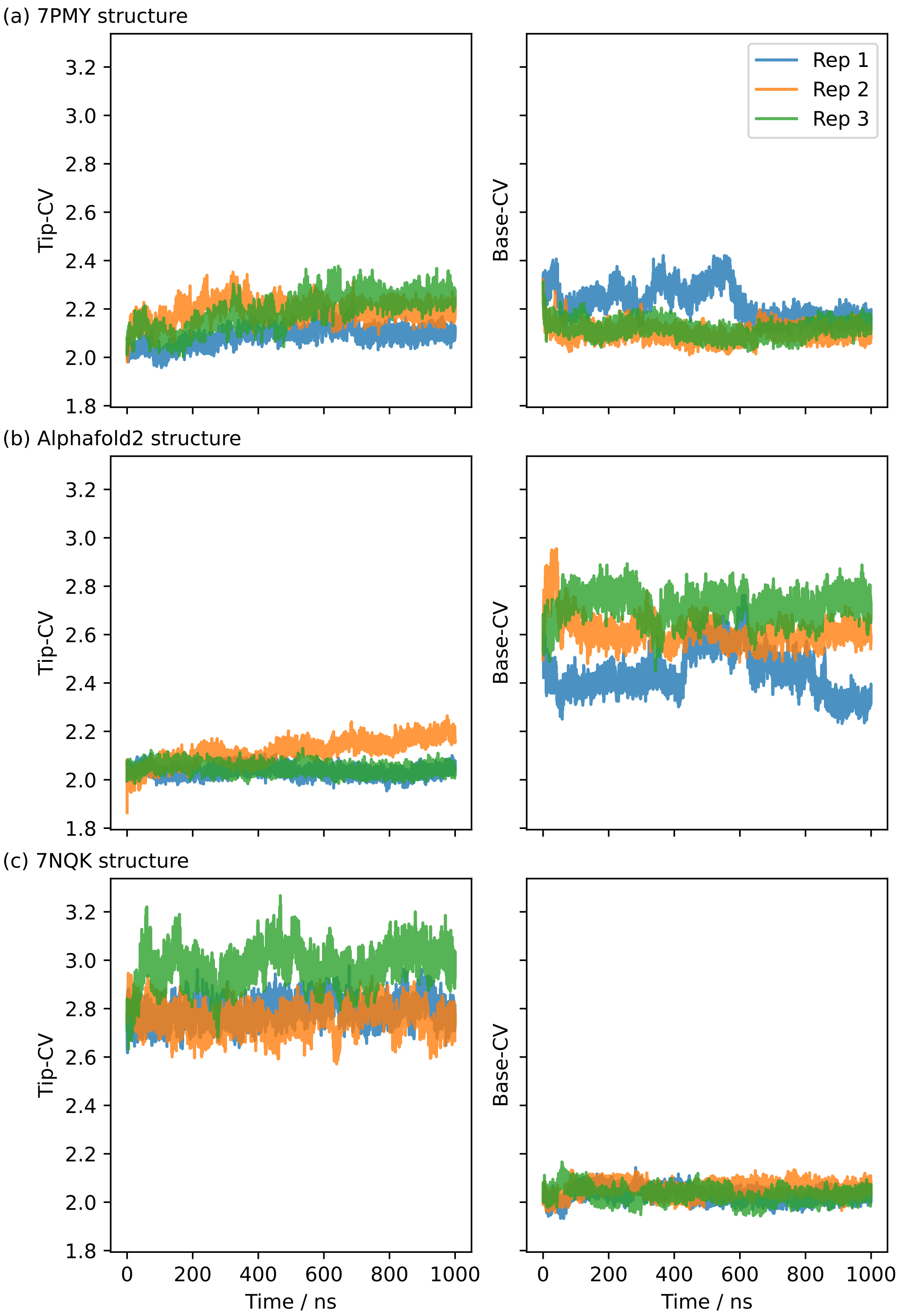


**Figure S1.** 1 μs-long MD simulations starting from CHARMM-GUI-embedded and equilibrated PepT2 structures. (a) The IF, partially OCC cryo-EM structure (7PMY) moves towards an occluded state via closure of the intracellular gate. However the extracellular gate partially opens in the process. Rep 1 also displays a partial helical unfolding near the intracellular gate (see provided coordinate files in the supplementary data). (b) Alphafold-based IF embeddings 1 and 3 explore a range of IF conformations while maintaining a stable extracellular gate, whereas rep 2 partially opens the extracellular gate. (c) The OF cryo-EM structure (7NQK) remains stable with a tight intracellular gate.

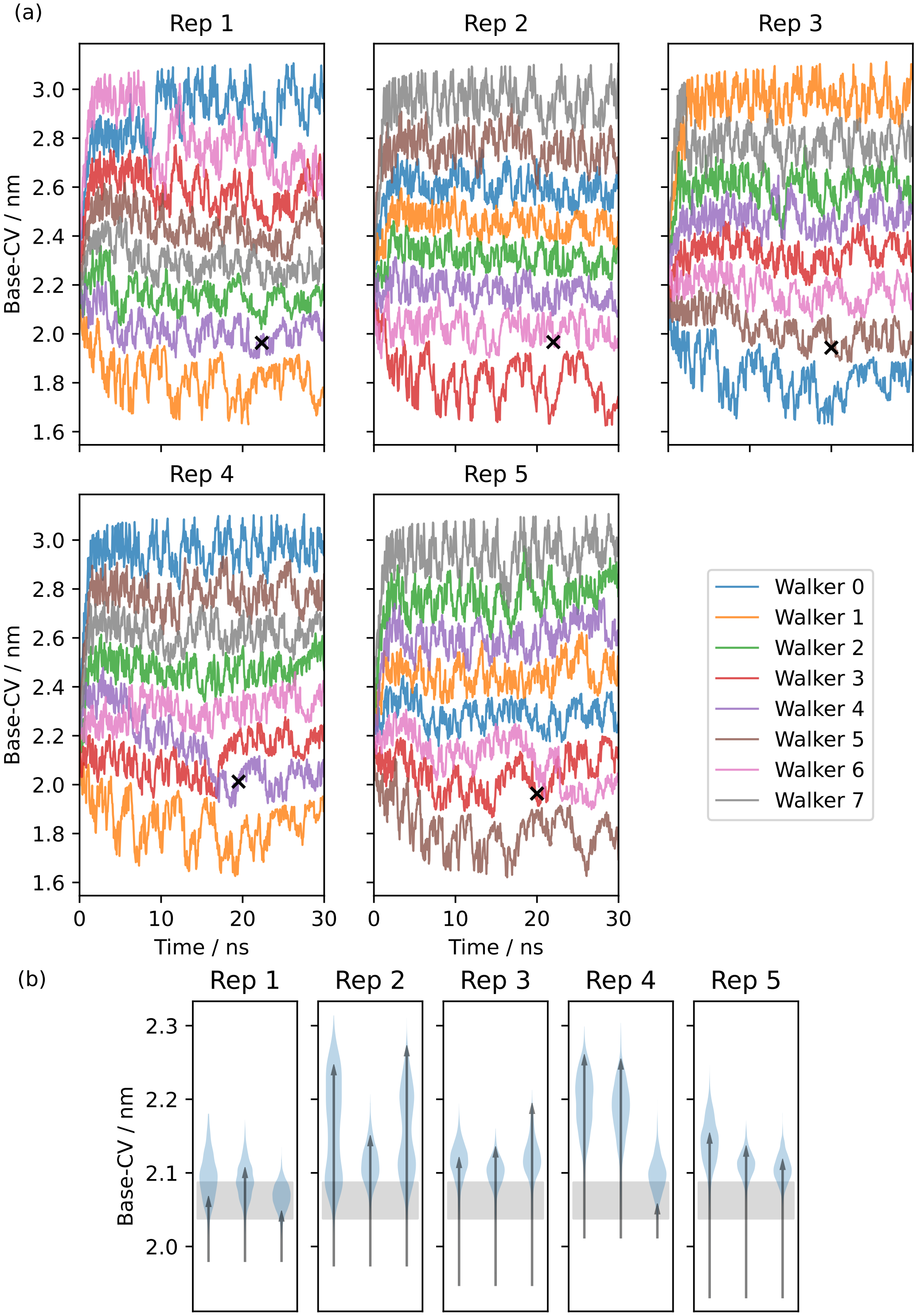


**Figure S2.** (a) Five replicates of multiple-walker metadynamics along the Base-CV, starting from the Alphafold IF equilibration rep 1. Black crosses indicate potential OCC states picked around a Base-CV value of 2 nm and at a timestep of around 20 ns. (b) Equilibrations (3 per metadynamics replciate) of the candidate OCC states in 100 ns unbiased MD. Histograms of trajectory projections onto the Base-CV are shown as violin plots, arrows link the first and final frames. The grey shaded area corresponds to the range of Base-CV values sampled in our OF simulation. Rep 1 is a stable OCC state, whereas the other replicates display partial intracellular-gate opening.

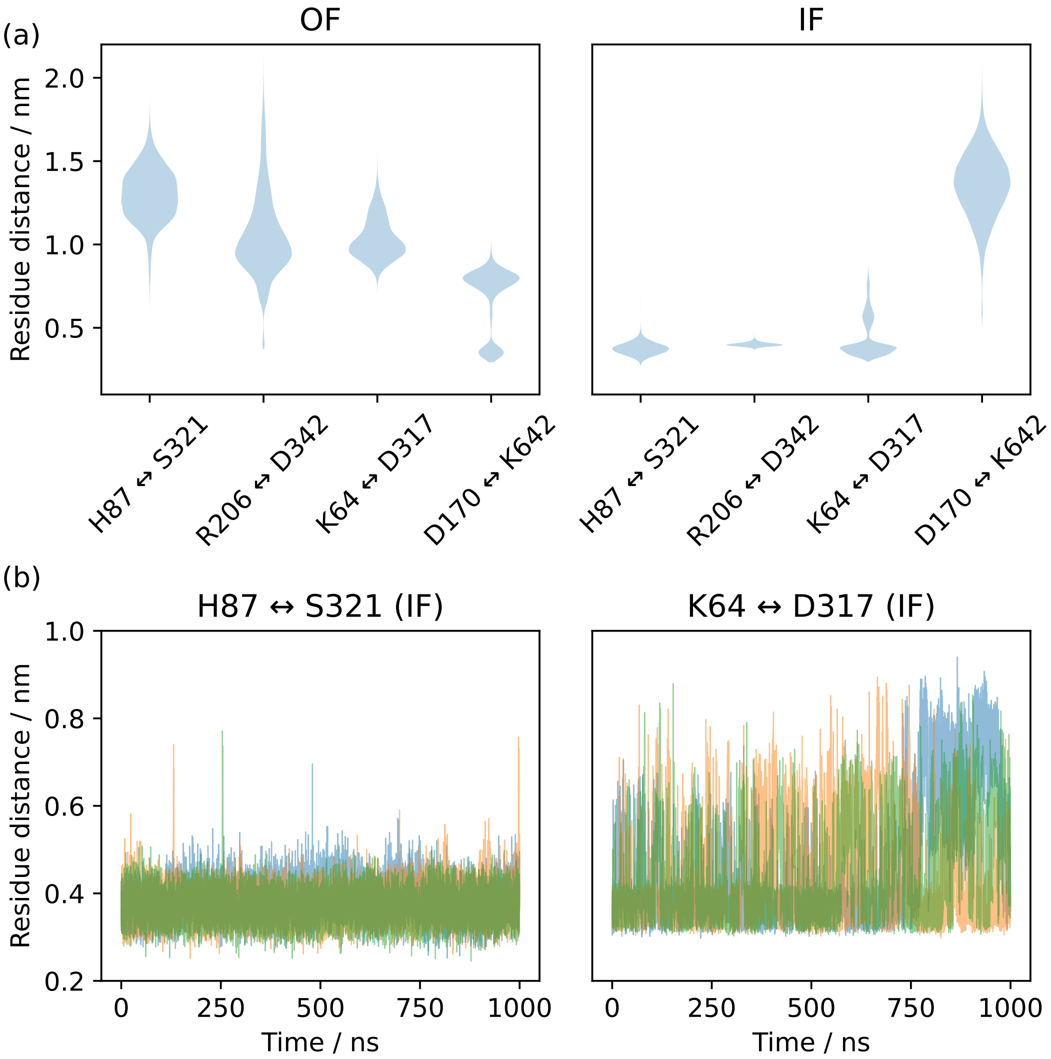


**Figure S3.** Inter-residue heavy atom (H87: NE2, S321: OG, R206: CZ, D342: CG, K64: NZ, D317: CG, D170: CG, K642: NZ) distances for several possible gating interactions. (a) Histograms from pooled triplicates of 1 µs MD as violin plots in the top row, starting from alphafold-derived IF and Cryo-EM derived OF conformations. H87↔S321 and R206↔D342 are always formed in the IF state but not in the OF state, while K64↔D317 and D170↔K462 show a preference for the IF and OF states respectively but are not formed in all trajectory frames. (b) Triplicate time series of the H87↔S321 and K64↔D317 interactions in the IF state, showing how the former is a tight interaction, while the latter is unstable and only transiently formed.

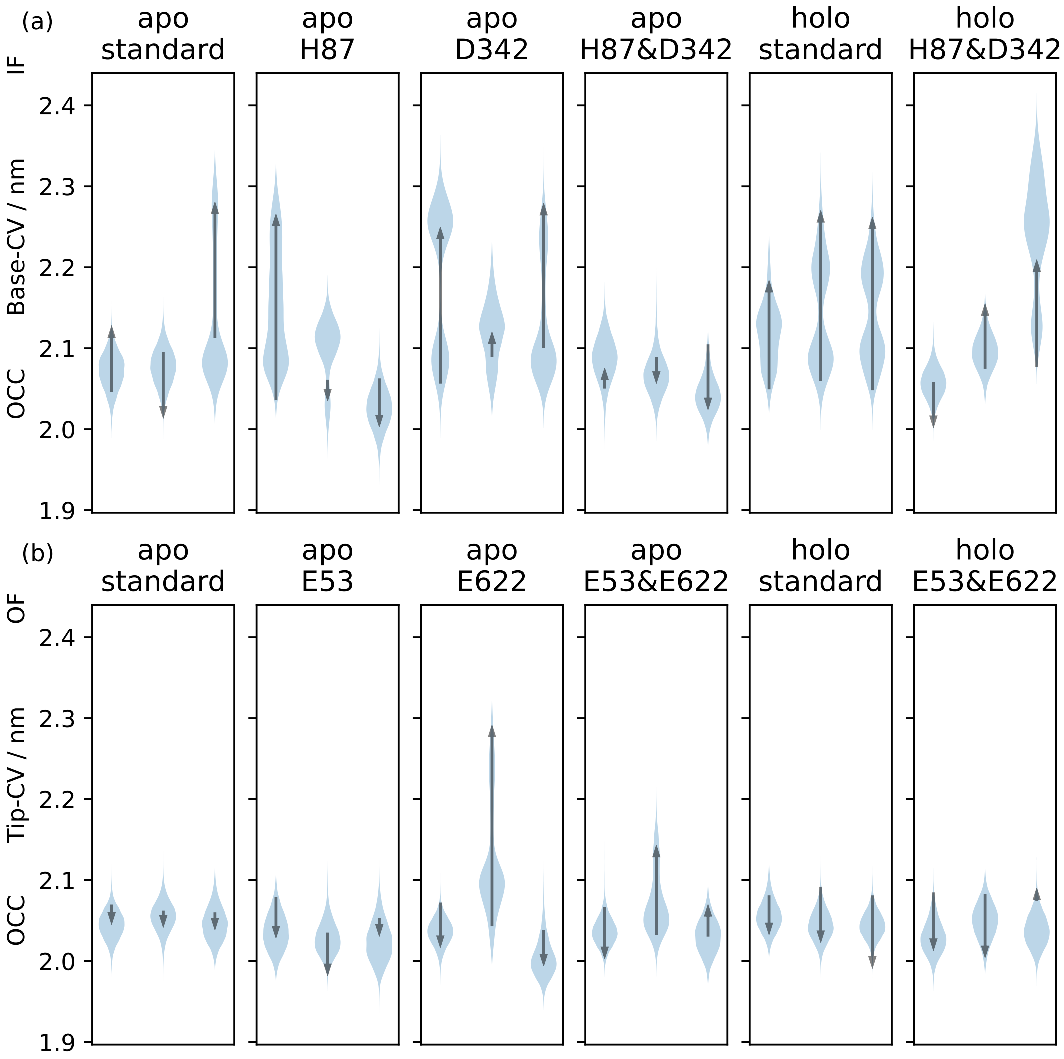


**Figure S4.** Triplicate 1 µs-MD simulations starting from OCC, showing the effects of different protonation and substrate binding states, projected onto the (a) Base-CV and (c) Tip-CV respectively. Violin plots are trajectory histograms, arrows link the CV values of the first and last frames. Intracellular gate flexibility is suppressed by conditions that favour extracellular gate opening and vice versa.

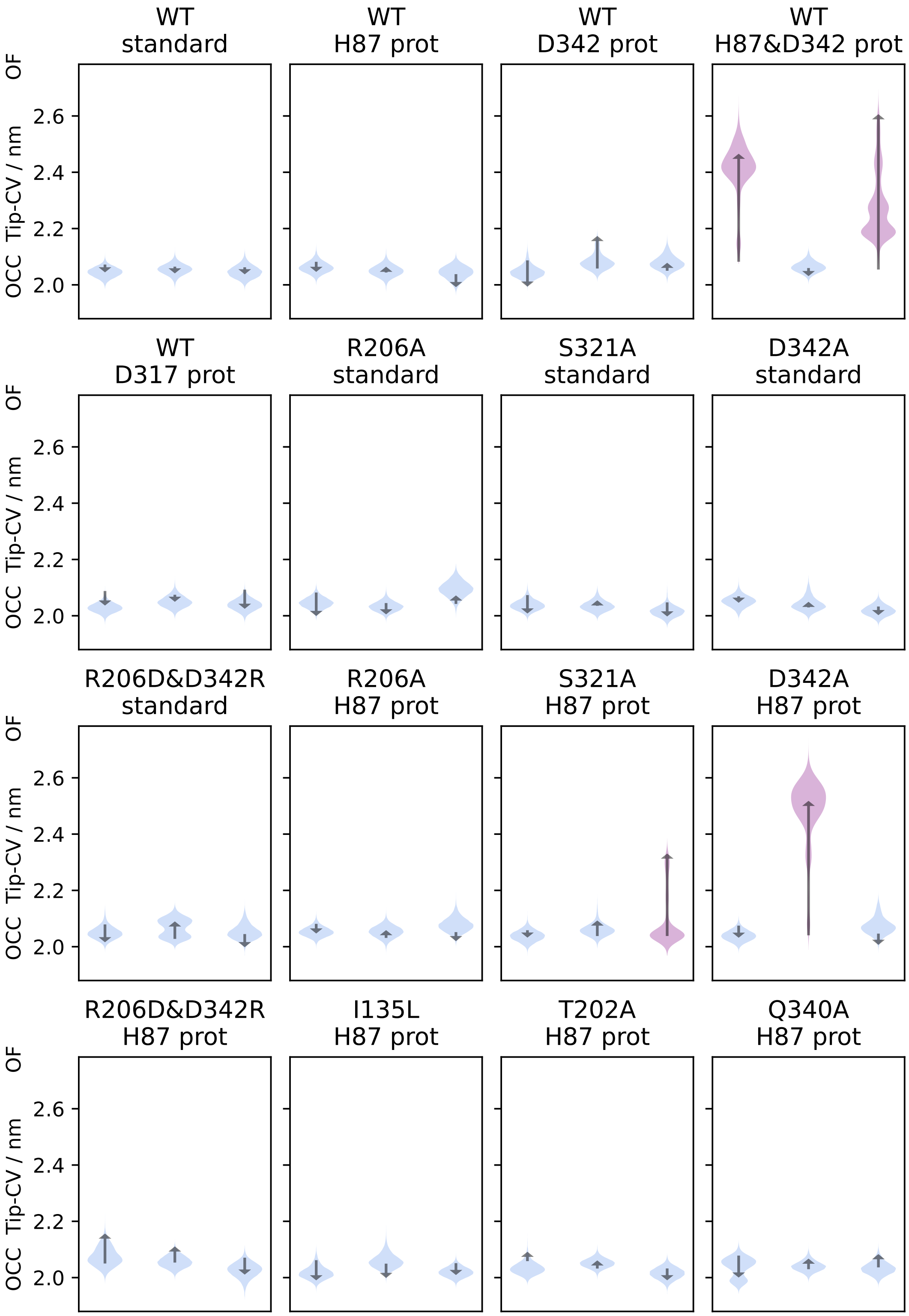


**Figure S5.** Triplicate 1 μs-MD simulations starting from the OCC state, showing the effects of different protonation states and mutations projected onto the tip-CV. Violin plots are trajectory histograms, arrows link the CV values of the first and last frames. Trajectories which displayed significant extracellular gate opening are highlighted in purple. Spontaneous extracellular gate opening requires H87 protonation, and the disruption of the R206↔D342 salt bridge also makes a significant contribution, either by mutation or protonation.

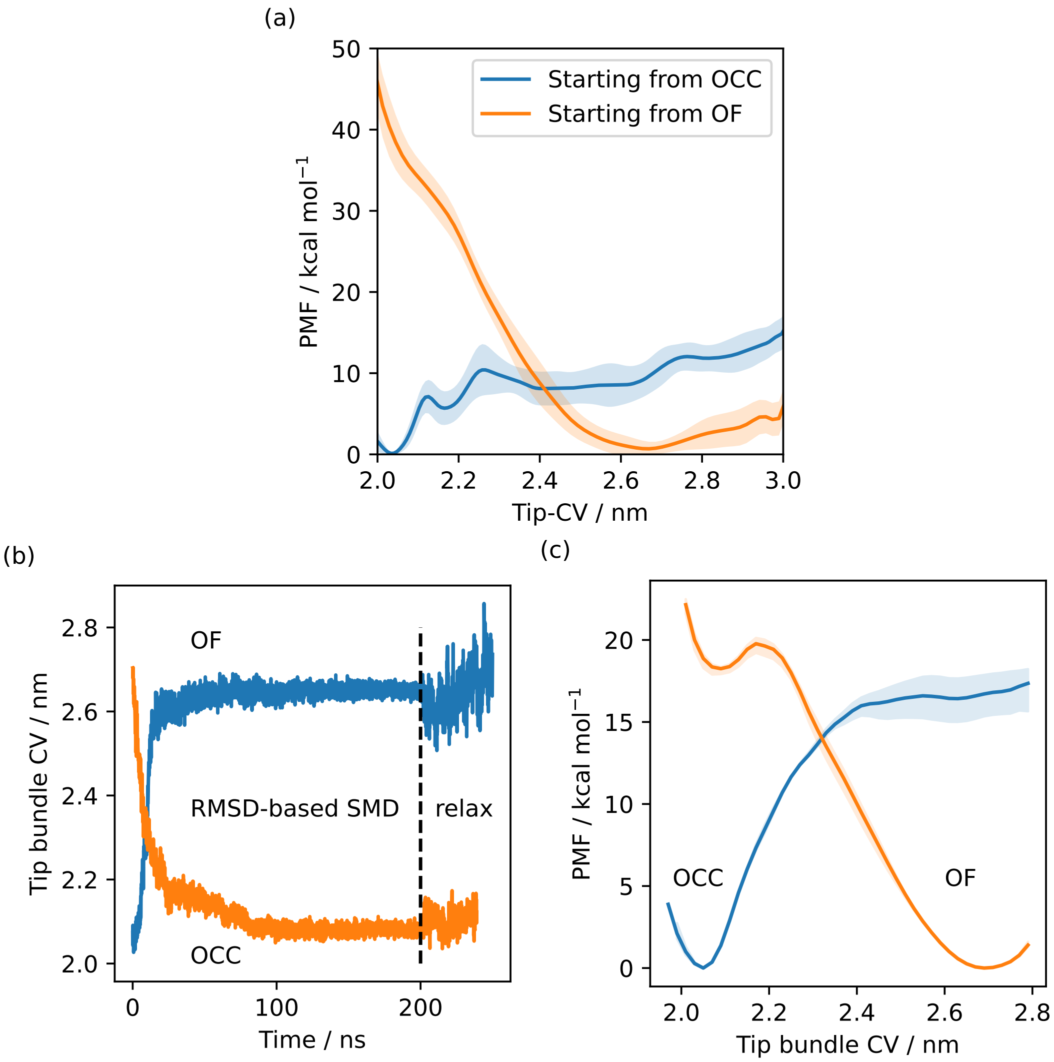


**Figure S6.** (a) Free energy profiles from metadynamics simulations (8 walkers) along the Tip-CV, starting simulations from the OCC (total sampling 1.7 µs) or OF state (total sampling 860 ns). Solid lines are the free energy estimates using the second half of the data only, shaded area is the standard deviation of free energy estimates with respect to sequential data chunks. The disparity between the curves indicates a significant hysteresis problem, favouring the initial state of the respective metadynamics run. (b) SMD runs between OF and OCC states, biasing the heavy-atom RMSD to the respective target state, shown as projections along the tip-CV. Metastable OCC and OF states are formed that remain stable in 50 ns unbiased MD. (c) REUS along the tip-CV with starting conformations picked to be equidistant in the CV from two SMD runs. Sampling was using 48 windows for a total of 4.4 µs (OCC→OF path) and 6.1 µs (OF→OCC path). Solid lines are PMFs calculated using all sampling, the shaded areas are error ranges obtained by omitting either the first 40% or the last 40% of sampling. The disparity between the curves indicates a significant hysteresis problem, favouring the initial state of the respective SMD path-generation run.

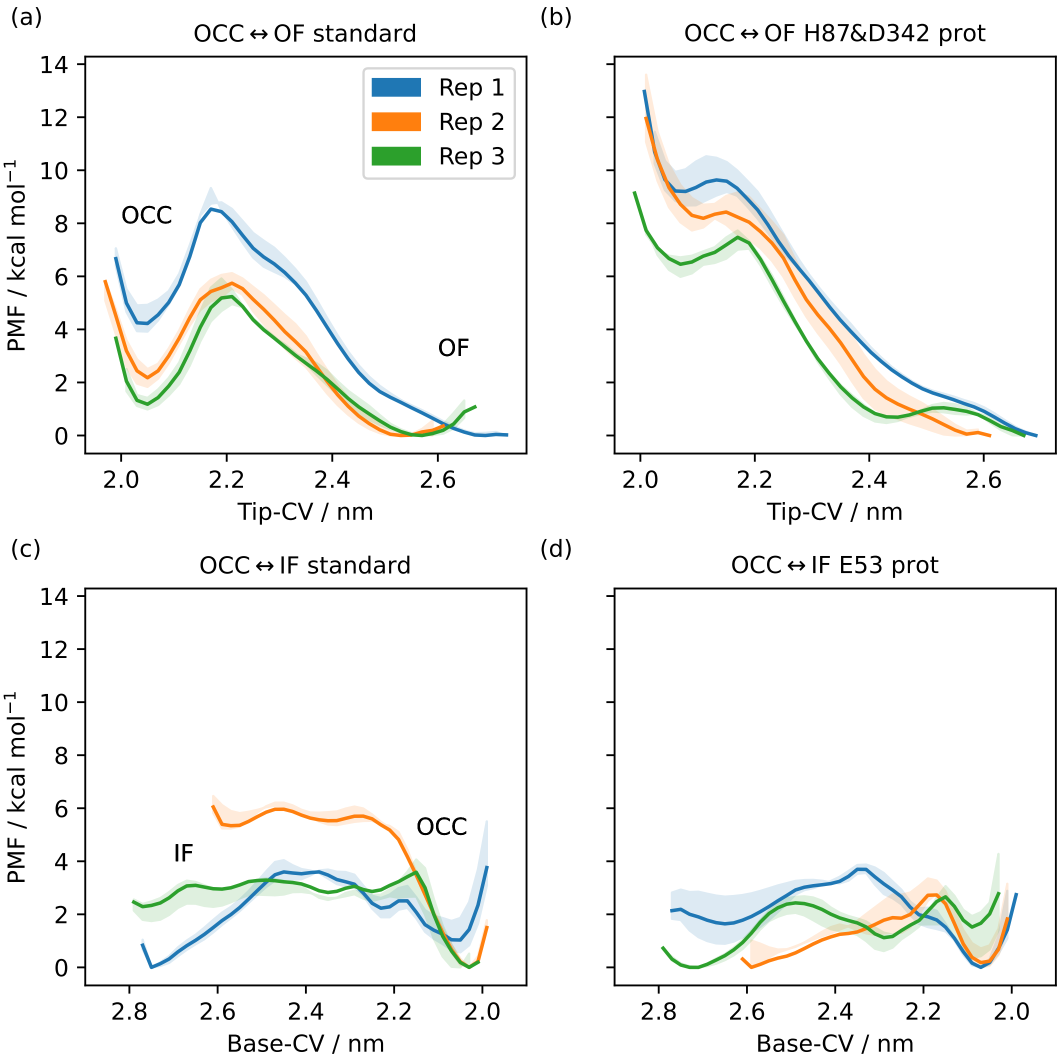


**Figure S7.** 1D-PMFs along the Tip-CV or Base-CV (as indicated), from REUS starting at MEMENTO intermediates. The central line is calculated using all sampling, whereas the shaded areas are enclosed by curves derived from omitting the first 40% or the last 40% of sampling, giving a sense of apparent convergence (while substantial inter-replicate differences remain). (a) OCC↔OF transition, standard protonation states. Distinct OCC and OF basins with separating barrier. (b) OCC↔OF transition, H87 and D342 protonated. The OCC basin and the separating barrier largely disappear. (c) OCC↔IF transition, standard protonation states. Distinct OCC basin and raised IF plateau. (d) OCC↔IF transition, E53 protontaed. The IF plateau and the barrier are lowered with respect to OCC.

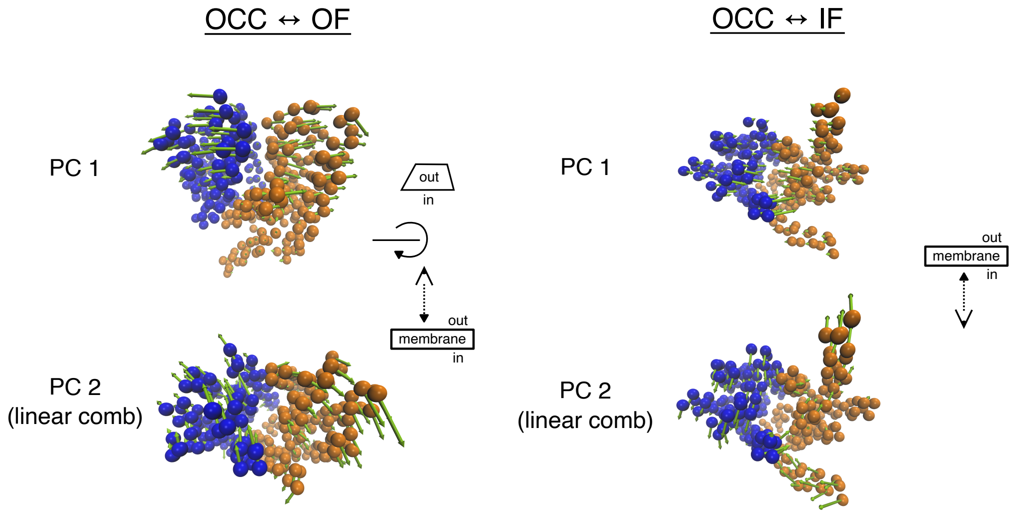


**Figure S8.** Illustration of the PCA-derived CVs for 2D-REUS. TM-helix CA-atoms are shown as spheres, coloured blue for the N-terminal bundle and orange for the C-terminal bundle, while arrows show magnitude and direction. For the OCC↔OF transition, PC 1 corresponds to the gating motion, while PC 2 is a cleft-sliding movement. For OCC↔IF, PC 1 corresponds to the gating motion, while PC 2 is a twisting movement. See supplementary videos 1–4 for animated versions of the same representations.

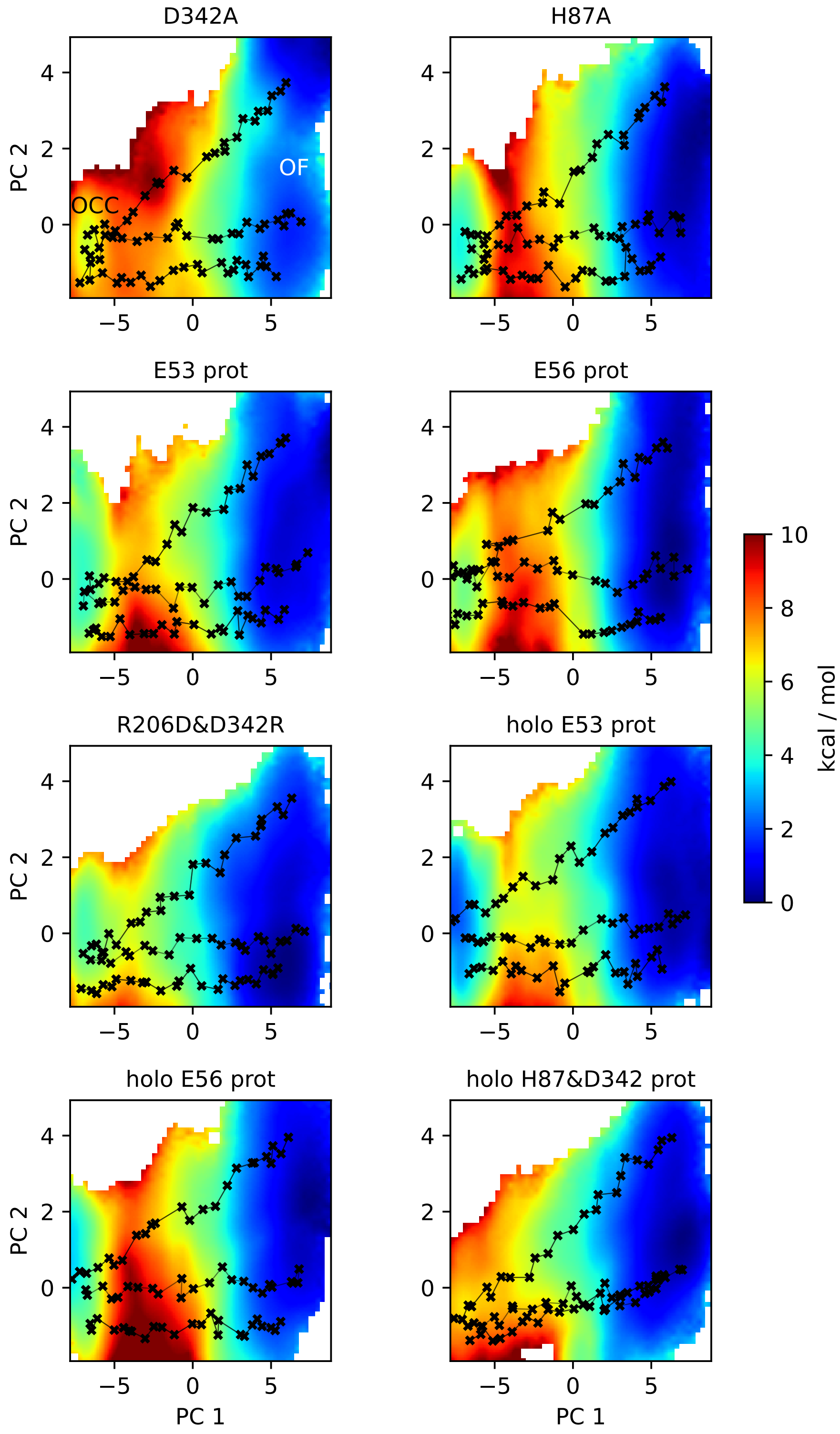


**Figure S9.** 2D-PMFs of the OCC↔OF transition from REUS with MEMENTO paths in additional protonation state and mutation conditions.

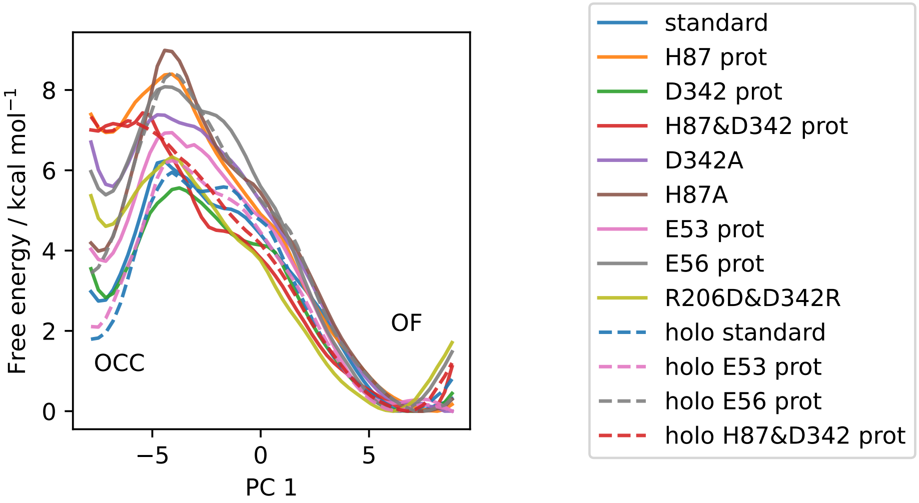


**Figure S10.** All available OCC↔OF 2D-PMFs, projected onto the first CV (PC 1). Solid lines are apo PMFs, dashed lines are ala-phe substrate-bound and color-matched to the respective apo PMF. Note that the individual PMFs are only determined by our REUS approach up to additive constants, and are shown aligned here at the OF state for convenience of comparison.

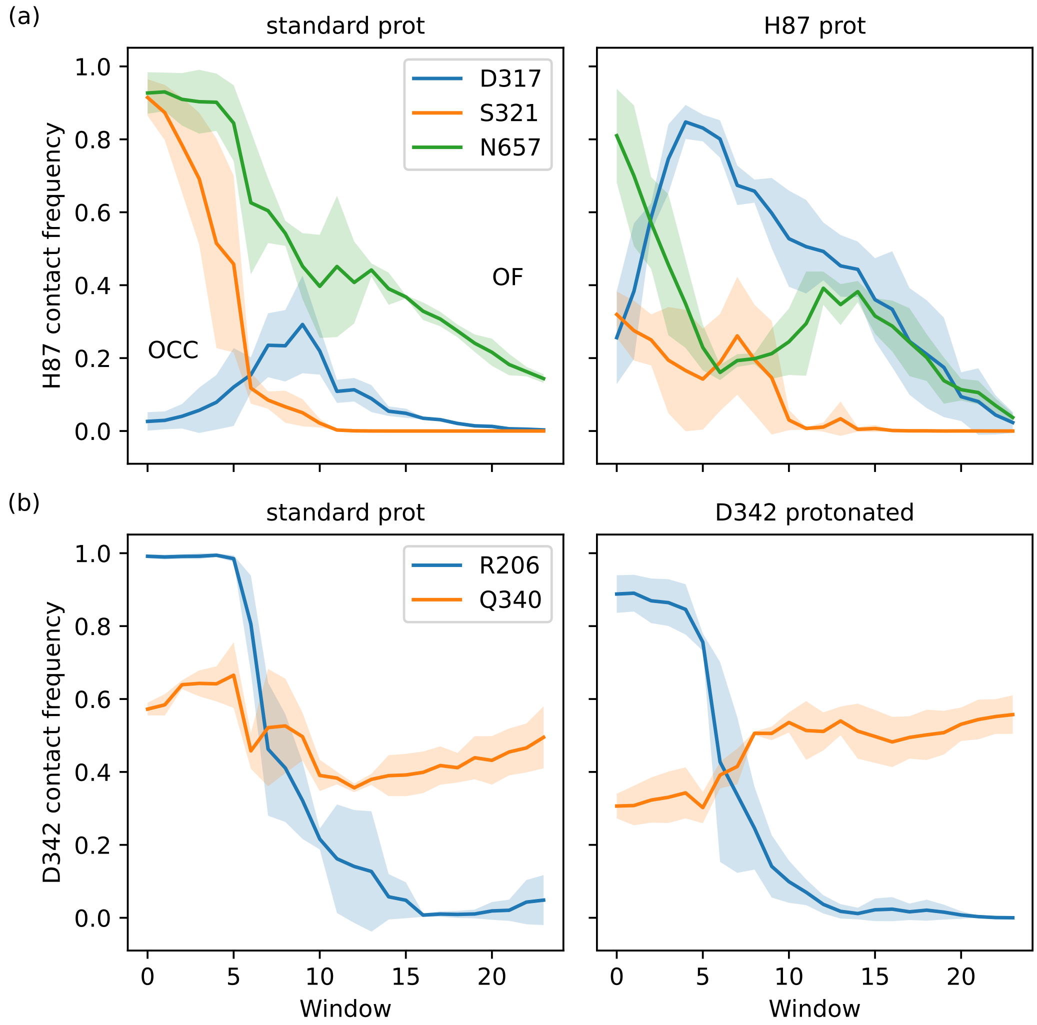


**Figure S11.** Interaction plots of the 2D-PMF trajectory data (high force constant windows only), calculated as frequencies of finding inter-residue heavy-atom distances smaller than 0.35 nm, shown as a line for the average across 3 replicates with shaded standard deviations. (a) H87 interactions with D317, S321 and N657. Protonation of H87 replaces the S321 interaction by an interaction with D317. (b) D342 interactions with R206 and Q340. The tight salt bridge D342–R206 is disrupted by D342 protonation, but the residues still interact in the OCC state via hydrogen bonds.

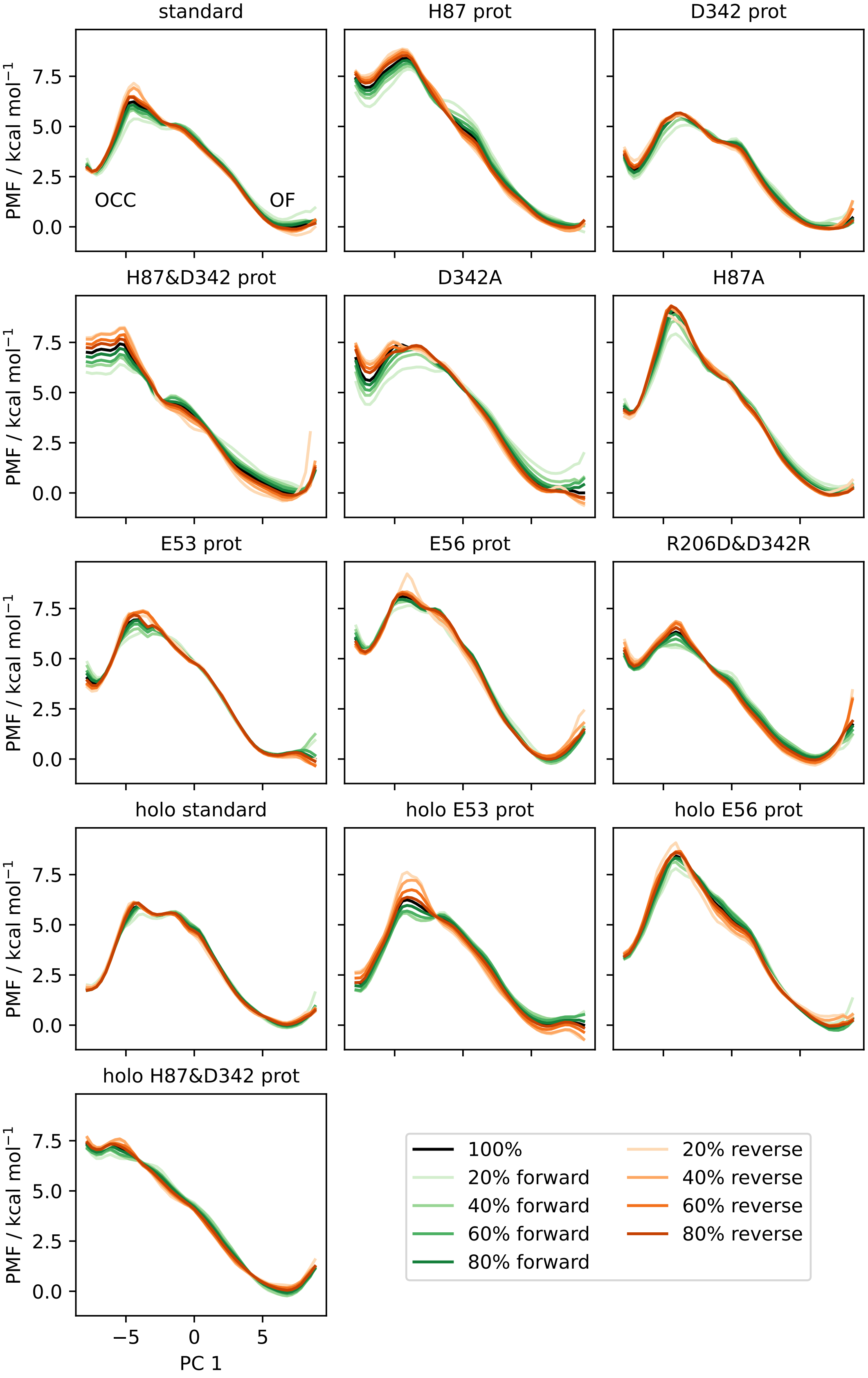


**Figure S12.** Convergence plots of all OCC↔OF 2D-PMFs, shown as projections onto PC 1 including successively (increasing saturation) more data points, starting from the first frame (green) or from the last frame in reverse (orange). The PMF using all data is shown in black.

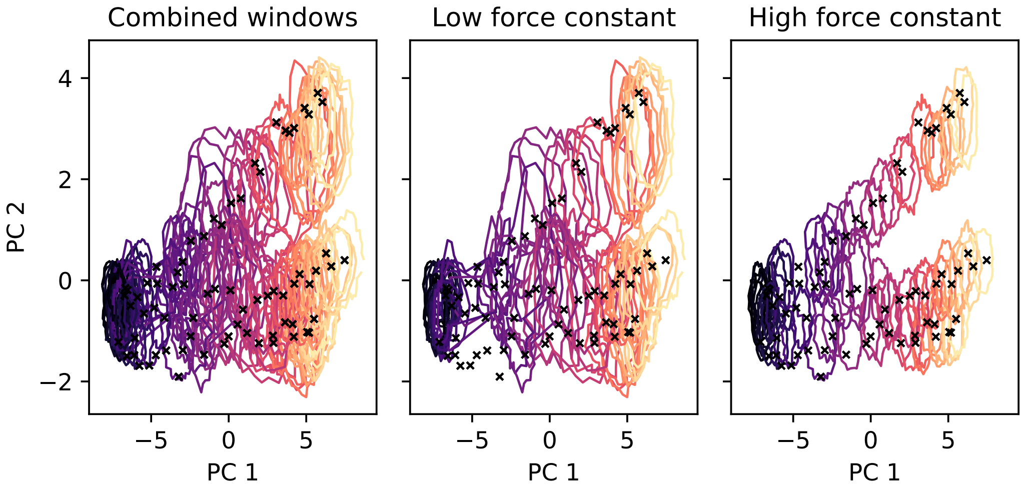


**Figure S13.** 2D-REUS histograms for the OCC↔OF standard protonation state 2D-PMF, drawn as contour lines at 30% of the maximal histogram height for each window (coloured by window, from purple at OCC to yellow at OF). Black crosses indicate the MEMENTO-derived REUS starting frames. Using the lower force constant (see Materials and Methods) gives good overlap in the basin regions while the transition region is undersampled. Using the higher force constant gives good overlap in the transition region while the basins are not sampled widely enough. Combining all windows results in good overlap across the 2D-CV space.

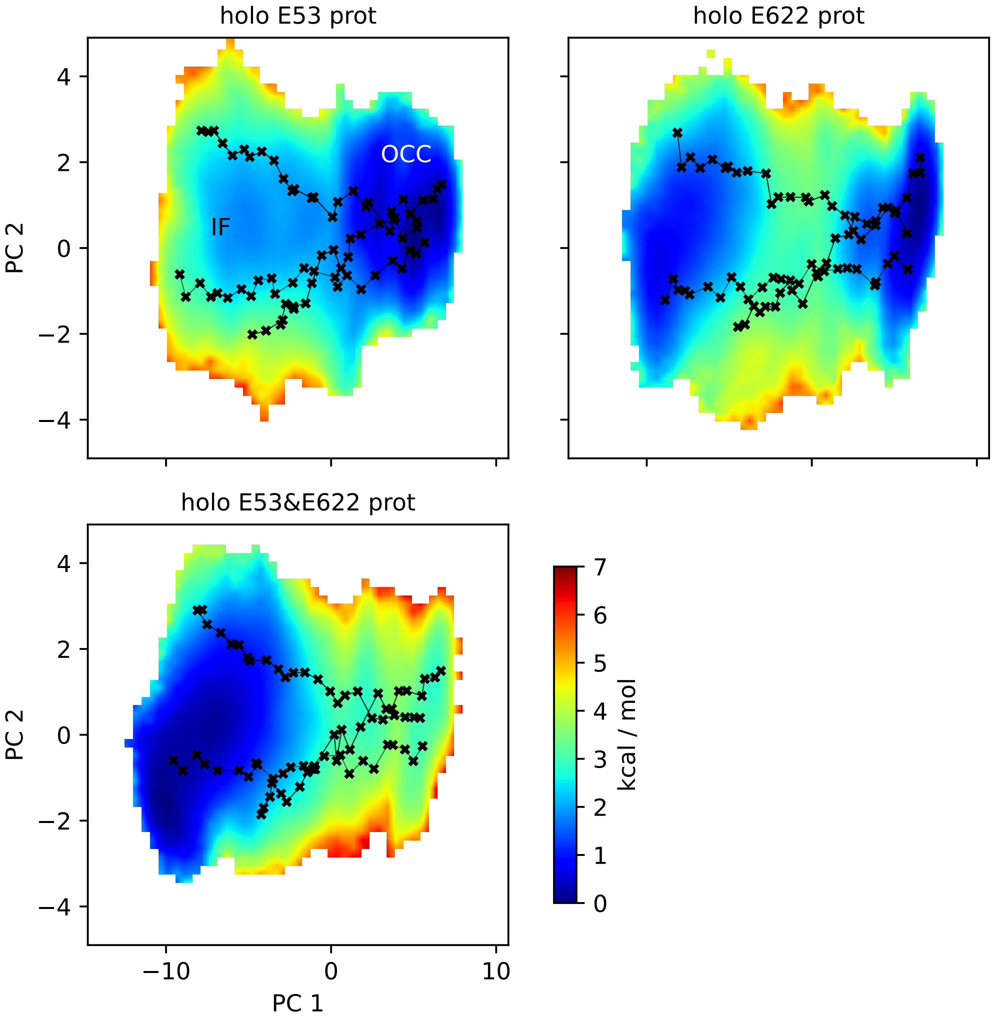


**Figure S14.** 2D-PMFs of the OCC↔IF transition from REUS with MEMENTO paths in additional substrate-bound protonation state conditions.

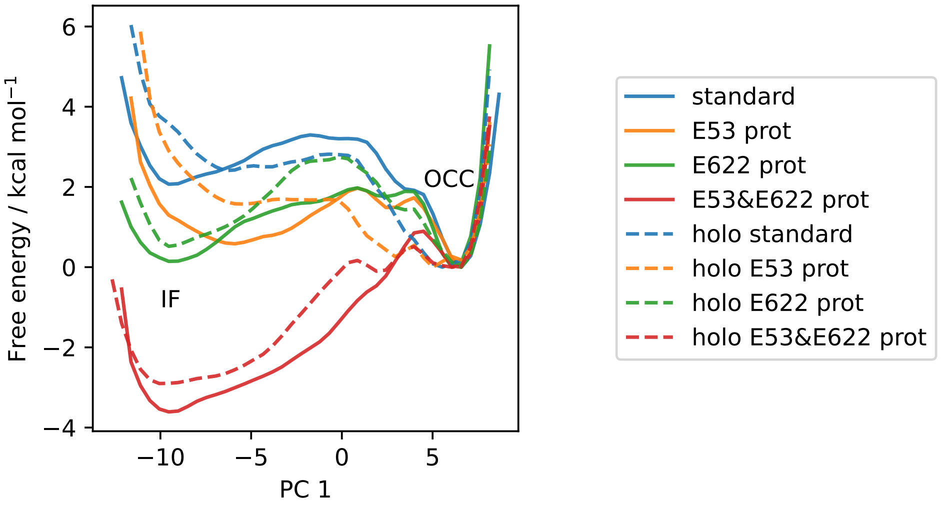


**Figure S15.** All available OCC↔IF 2D-PMFs, projected onto the first CV (PC 1). Solid lines are apo PMFs, dashed lines are ala-phe substrate-bound and color-matched to the respective apo PMF. Note that the individual PMFs are only determined by our REUS approach up to additive constants, and are shown aligned here at the IF state for convenience of comparison.

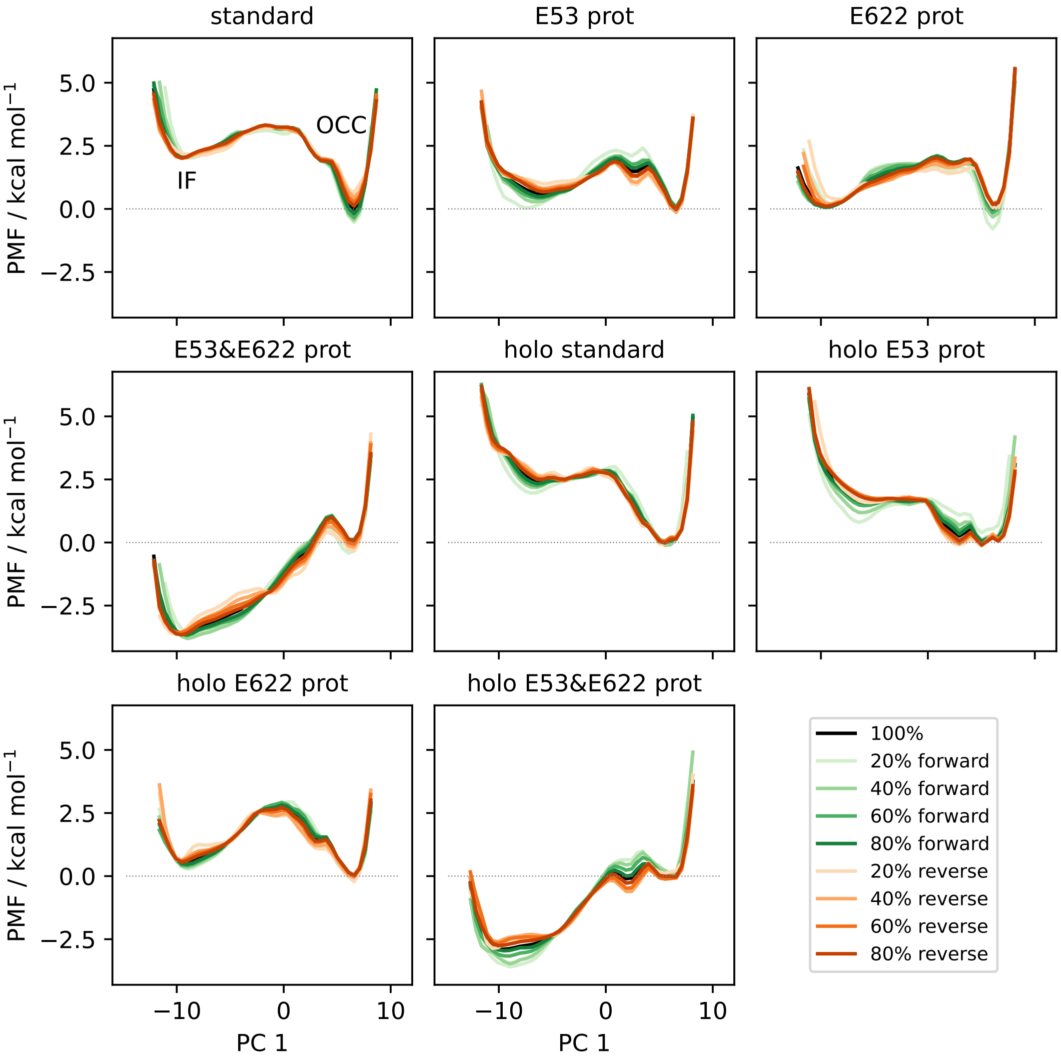


**Figure S16.** Convergence plots of all OCC↔IF 2D-PMFs, shown as projections onto PC 1 including successively (increasing saturation) more data points starting from the first frame (green) or from the last frame in reverse (orange). The PMF using all data is shown in black.

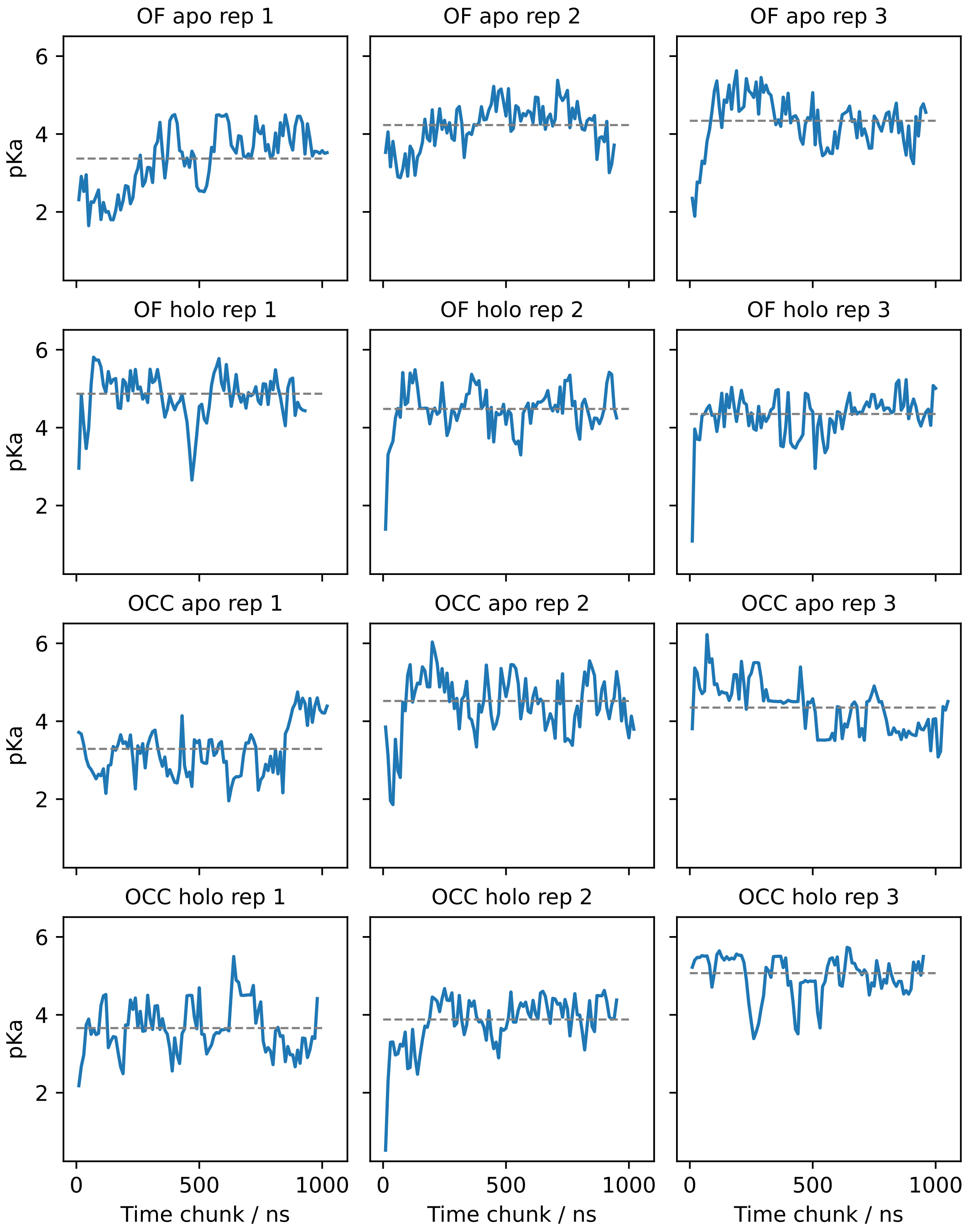


**Figure S17.** E53 pKa values estimated for (separate) successive chunks of 80 ns (10 ns per pH window) of CpHMD via fitting the Hill equation. Grey lines indicate the pKa resulting from fitting the total simulation data. The pKa values appear highly dynamic and superimpose several relaxation timescales. Convergence seems to be achieved in most replicates but is not straightforward to assess.

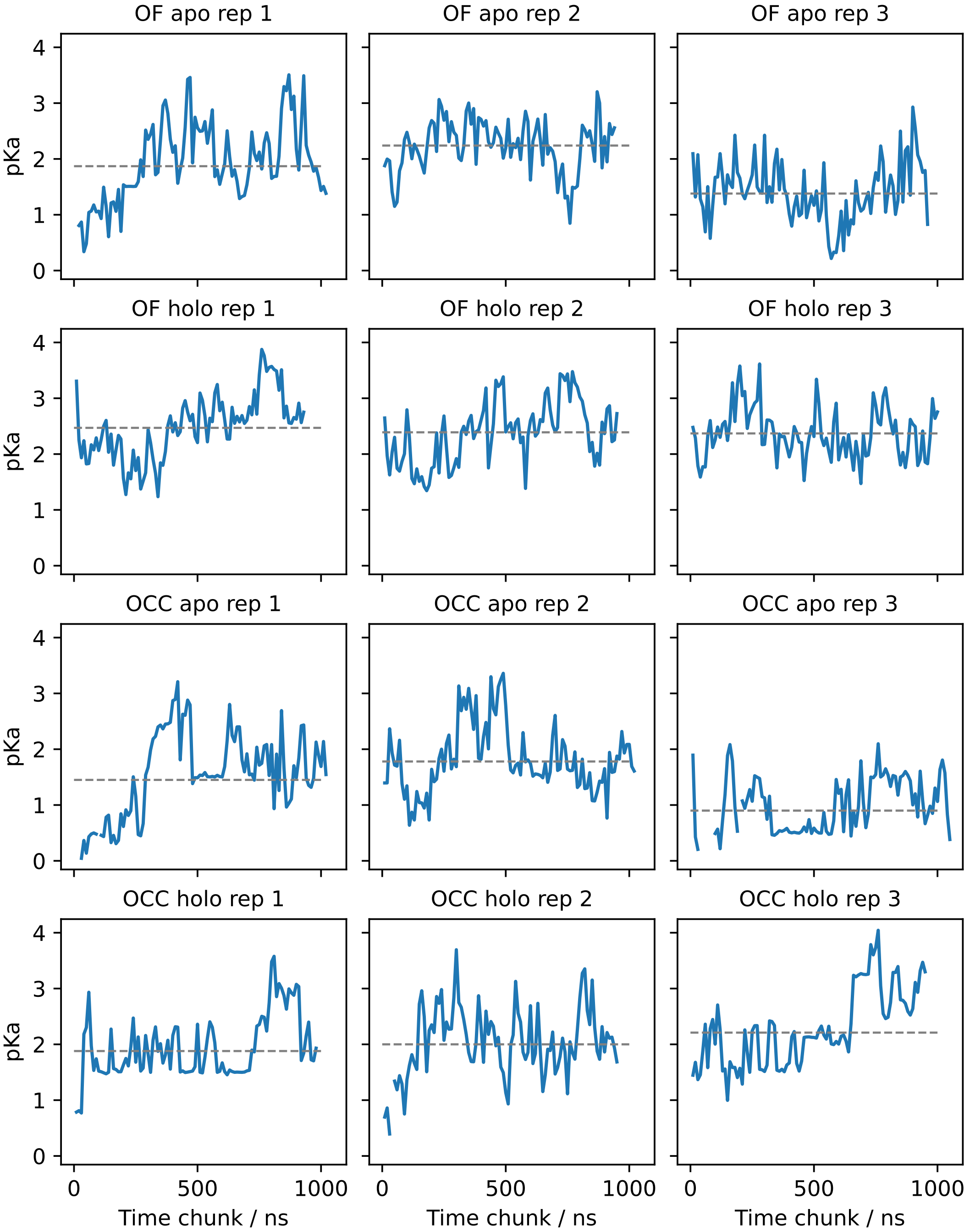


**Figure S18.** E56 pKa values estimated for (separate) successive chunks of 80 ns (10 ns per pH window) of CpHMD via fitting the Hill equation. Grey lines indicate the pKa resulting from fitting the total simulation data. The pKa values appear highly dynamic and superimpose several relaxation timescales. Convergence seems to be achieved in most replicates but is not straightforward to assess.

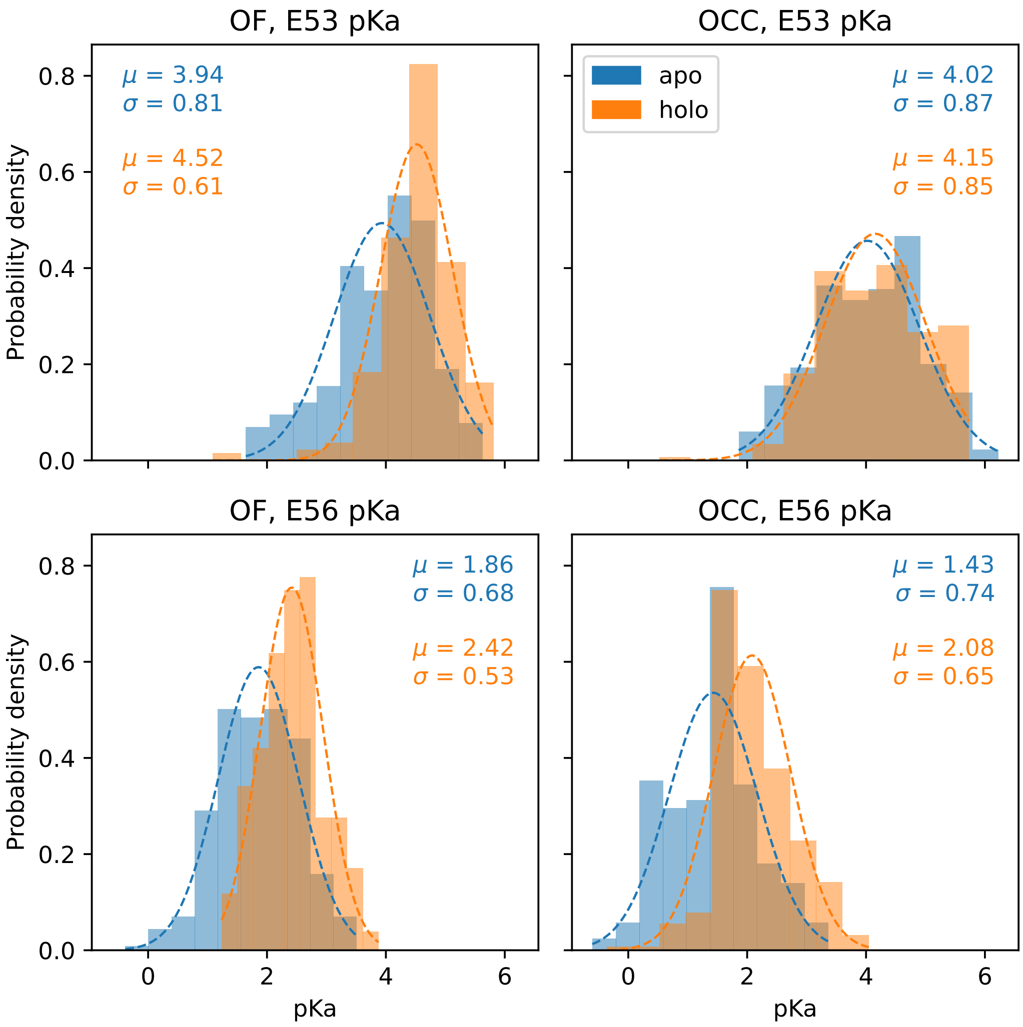


**Figure S19.** Histograms of the pKa values estimated from chunks of 80 ns (10 ns per pH window) of CpHMD, pooled for all trajectories of a given condition. Apo conditions are shown in blue, holo (ala-phe bound) are shown in orange. Gaussian fits are indicated by dashed lines and the resulting fit parameters are given in the respective color for each histogram.

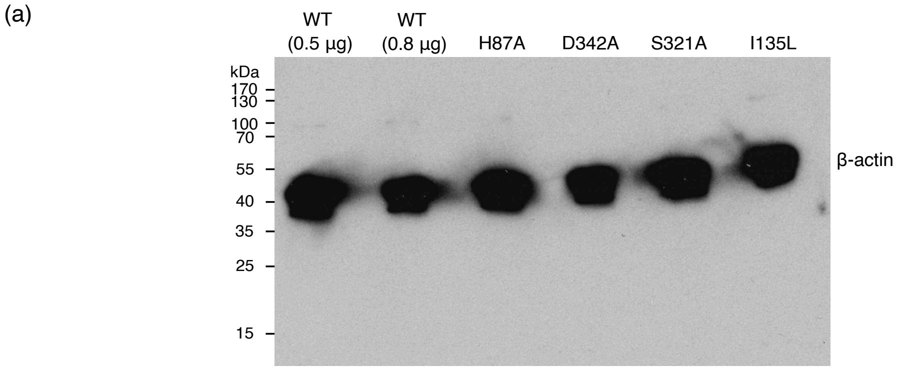


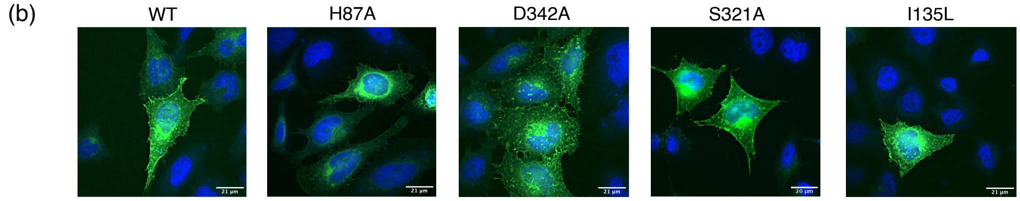


**Figure S20.** (a) Loading control of the Western blot shown in figure 7b, using an antibody against $\beta$-actin, showing even loading of the gel. (b) Fluorescence microscopy images, overlaying GFP-labelled PepT2 (green) with DAPI-labelled DNA (blue). Membrane expression is qualitatively shown for WT and all mutants by the thin cell outline of GFP.
